# *Bering:* joint cell segmentation and annotation for spatial transcriptomics with transferred graph embeddings

**DOI:** 10.1101/2023.09.19.558548

**Authors:** Kang Jin, Zuobai Zhang, Ke Zhang, Francesca Viggiani, Claire Callahan, Jian Tang, Bruce J. Aronow, Jian Shu

## Abstract

Single-cell spatial transcriptomics such as *in-situ* hybridization or sequencing technologies can provide subcellular resolution that enables the identification of individual cell identities, locations, and a deep understanding of subcellular mechanisms. However, accurate segmentation and annotation that allows individual cell boundaries to be determined remains a major challenge that limits all the above and downstream insights. Current machine learning methods heavily rely on nuclei or cell body staining, resulting in the significant loss of both transcriptome depth and the limited ability to learn latent representations of spatial colocalization relationships. Here, we propose ***Bering***, a graph deep learning model that leverages transcript colocalization relationships for joint noise-aware cell segmentation and molecular annotation in 2D and 3D spatial transcriptomics data. Graph embeddings for the cell annotation are transferred as a component of multi-modal input for cell segmentation, which is employed to enrich gene relationships throughout the process. To evaluate performance, we benchmarked *Bering* with state-of-the-art methods and observed significant improvement in cell segmentation accuracies and numbers of detected transcripts across various spatial technologies and tissues. To streamline segmentation processes, we constructed expansive pre-trained models, which yield high segmentation accuracy in new data through transfer learning and self-distillation, demonstrating the generalizability of *Bering*.

## Introduction

In recent years, there has been a significant advancement in single-cell spatial transcriptomics technologies, providing powerful tools to study transcript localization and cellular processes at a high resolution and scale^1,2^. These innovative technologies comprise *in-situ* hybridization methods, such as MERFISH and SeqFISH^3,4^, and *in-situ* sequencing approaches such as STARmap, pciSeq^5–7^. Besides, Several commercially available technologies, such as MERSCOPE, Nanostring CosMx, and 10x Xneium, have been developed and made the spatial techniques more accessible to researchers^8–10^. Although single-cell spatial transcriptomics technologies were initially limited by the need for multiple rounds of tissue staining, resulting in lower throughput compared to Next-Generation-Sequencing (NGS) technologies, this constraint is outweighed by the ability to achieve high resolutions of hundreds of nanometers, reaching subcellular resolutions^11^. This attribute allows for extensive exploration of cellular processes and interactions at high spatial resolution. Moreover, cutting-edge image-based spatial transcriptomics technologies, including seqFISH^+^ and STARmap PLUS, have achieved significant advancements in gene throughput, reaching several thousand or even tens of thousands^4,5^. As a cutting-edge technology, single-cell spatial transcriptomics renders rich information on transcripts’ spatial distribution at remarkably high resolutions, while staining images of the data offer valuable insights into cell morphological characteristics.

Despite the vast potential of single-cell spatial transcriptomics technologies, analyzing the resulting data presents several complex computational challenges. One of the most significant obstacles is cell segmentation, which is critical for accurately delineating individual cells within a tissue sample in spatial data. Pioneering deep learning approaches, such as Cellpose and JSTA^12,13^, have proven effective for cell segmentation tasks using nuclei staining. However, there are two challenges associated with this strategy. Firstly, a significant number of transcripts are present in both the nuclei and cytoplasm, making them difficult to fully capture through nuclei staining alone^6^. Additionally, this strategy is unable to capture transcript spatial patterns or their colocalization, missing out valuable insights into cell compartments and structures^14^. For instance, transcription factors, such as *OCT4*, and histone genes, are predominantly found in nuclei, whereas cytoskeletal protein genes such as *TLN1* are more commonly observed in the cytoplasm and membrane^15^. In light of this, some methods have sought to leverage spatial distributions of transcripts for cell segmentation in spatial transcriptomics data, such as ClusterMap and Baysor^16,17^. However, it remains challenging for these statistical methods to efficiently learn the latent representation of transcript colocalization relationships within such high-dimensional spatial data. An innovative approach taken by SCS involves the utilization of transformer models on integrated imaging and transcript data to enhance cell segmentation accuracy. Nevertheless, one of the core steps of SCS relies on the identification of cell nuclei based on nuclei staining, which is often observed to be incomplete in its coverage of cells^18^. Consequently, this strategy may lead to a significant loss of cells and transcripts.

In the face of these challenges, we embarked on a comprehensive exploration of the multi-modal data in single-cell spatial transcriptomics. We found significant loss of transcripts during segmentation in strategies that sorely rely on staining images, and highlighted gene patterns that are indicative of cell types and boundaries from transcript colocalization data. To tackle challenges in segmentation and build upon discoveries of transcript colocalization, we introduce a computational approach that utilizes a graph neural network to harness transcript colocalization relationships for cell-type annotation. Notably, the learned transcript representations are transferred to the segmentation task as a component of multi-modal learning input, circumventing the limitations of single-modal learning. Innovatively, we approached the segmentation task as the edge prediction task to fully leverage transcript colocalization relationships and achieve a finer level of segmentation compared to conventional pixel-level segmentation methods. We have successfully applied this method to various tissue types and technologies, whether image-free or image-dependent, and have showcased its superior performance in accurately identifying precise cells in 2D and 3D thick tissues. Additionally, we demonstrate the generalizability of our approach by transferring the pre-trained model to a completely new dataset and achieving accurate cell segmentation results through self-distillation, highlighting the potential of broad applications across various tissues, volumes, and technologies.

## Results

### Spatial transcriptomics data modalities for segmentation analysis

Multiple types of staining images, such as DAPI, poly-A, and membrane staining, have been generated across spatial datasets and technologies for cellular morphological detection and cell segmentation (Fig. 1a, Supplementary Fig. 1). Among them, DAPI is the most widely used staining image for cell segmentation. However, when we projected transcripts onto the paired DAPI staining, we observed that a significant number of spots were not overlapped with strong DAPI signals, varying from 30% to 70% across samples and datasets (Supplementary Fig. 2), which can lead to loss of information during segmentation. While membrane staining can provide rich information for segmentation^17^, its inadequate and imbalanced imaging signals across different cell types could cause biased segmentation and loss of information (Supplementary Fig. 3).

**Figure 1.**
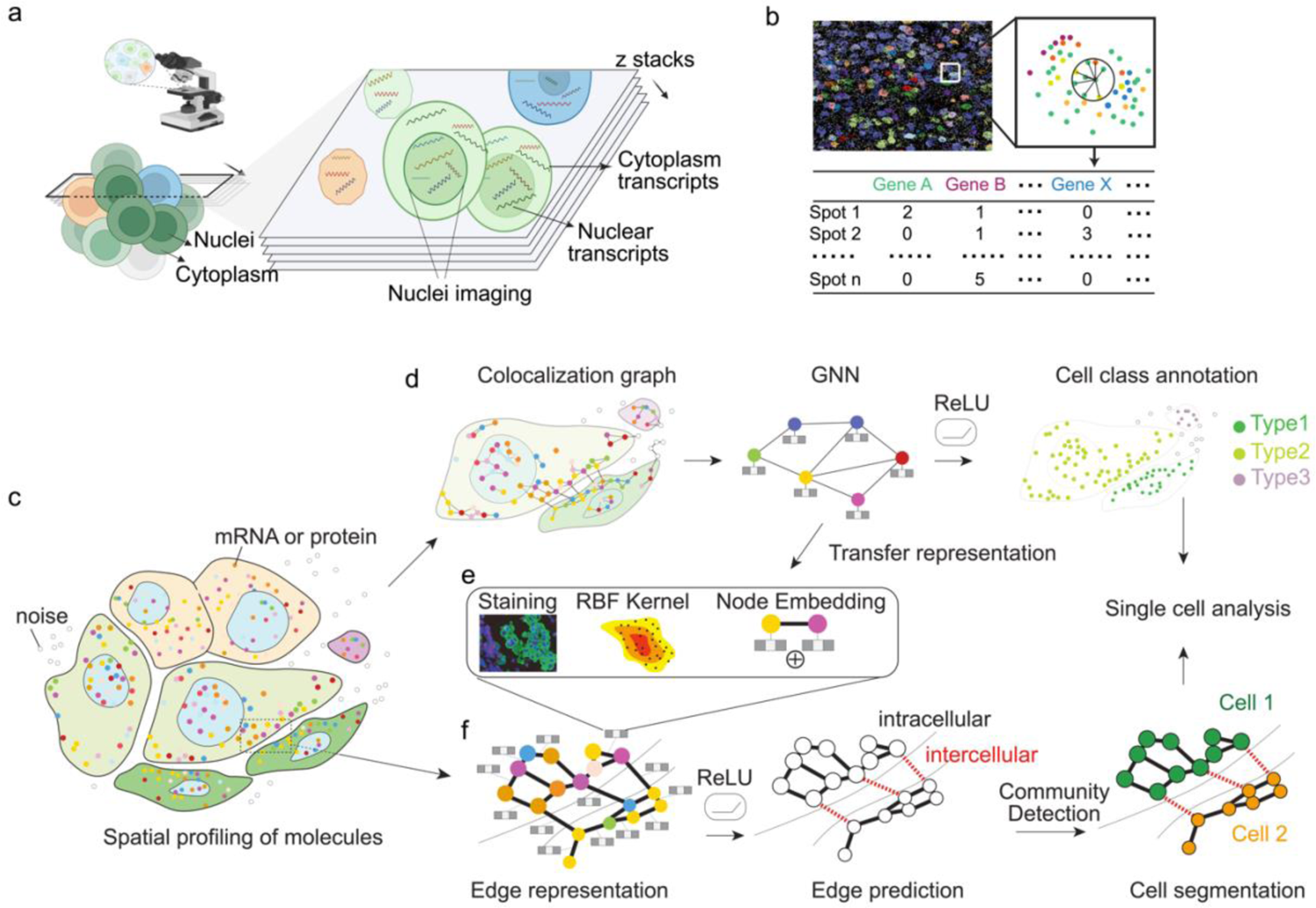
The overview of Bering model for cell segmentation. (a) An animation illustrating image-based spatial transcriptomics approaches. Multiple slices in the z-axis are generated and transcripts from nuclei and cytoplasm are detected in each slice. Additionally, staining images, including the nuclei image, are captured. (b) The concept of Neighborhood Gene Components (NGCs) is introduced for transcripts in image-based spatial transcriptomics data. NGCs are defined as count matrices, where each value in the matrix corresponds to the number of detected transcripts for each gene in the neighborhood of the query transcript. (c-f) Overview of the Bering model. (c) The animation of spatial profiling of molecules in single-cell spatial omics data. Colored spots indicate mRNA or proteins, while hollow spots represent noise. Different colors of cells indicate distinct cell types, with the nucleus shown in blue. (d) Construction of spatial colocalization neighborhoods using k-nearest graphs, with graph convolutional neural networks (GCN) trained for cell type prediction on the node level (transcript or other molecules). (e) Input for the edge prediction task, including required node embeddings transferred from the node classification task, along with auxiliary input of staining images and distance information calculated from the 2/3-dimensional (2/3D) coordinates. (f) Cell segmentation task defined as an edge prediction task, utilizing multimodal input of edge embeddings (from e) to train neural networks. The prediction outcome is binary classification, indicating whether the edges connect intercellular or intracellular spots. Molecular connectivity graphs are then constructed, and community detection algorithms such as Leiden Clustering are employed to identify cell borders. Results from the node classification task and segmentation task are combined to obtain single cells with annotations for downstream analysis. More details can be found in Supplementary Figure. 6.

To gain a holistic understanding of the single-cell spatial omics data, we delved into transcript profiles, and revealed their patterns of cell compartments and subcellular structures^11^. We utilized a factor analysis model^19^ in non-small cell lung cancer (NSCLC) and identified three distinct subcellular gene patterns in tumor cells, including nuclear genes (factor 2) and peripheral genes (factor 3) (Supplementary Fig. 4a-c). Nucleus-specific genes, such as *MALAT1* and *NEAT1*, exhibit a high enrichment within the nuclei region. In contrast, genes involved in kinase phosphatase activity, such as *DUSP5*, exhibit a notable enrichment within the cytoplasm of cells (Supplementary Fig. 4d-e), providing compelling evidence of a subcellular pattern as indicated by the spatial distribution of transcripts. To further understand the underlying information of transcript physical colocalization, we constructed neighborhood gene components (NGCs) by grouping the nearest transcripts together (Fig. 1b) and examined their latent dimensions using Uniform Manifold Approximation and Projection (UMAP) (see Methods). Remarkably, the distributions of NGCs on UMAP aligned closely with cell type and compartment annotations, highlighting the highly indicative nature of transcript neighbors in relation to cell-type-specific subcellular structures and boundaries (Supplementary Fig. 5).

### Bering overview

To effectively tackle the challenges mentioned above and fully capitalize on the information embedded in transcript distributions within spatial transcriptomics data, we have developed Bering, a novel approach that combines graph neural network (GNN) and transfer learning for joint cell segmentation and annotation (Fig. 1c-f, Supplementary Fig. 6). Drawing inspiration from the insights gleaned through the analysis of transcript colocalization data, Bering begins by harnessing the colocalization graph as its input. Subsequently, we employ GNN to predict both noises and cell types associated with transcripts, during which the model acquires the ability to encode representations for transcripts, leveraging the intricate information related to cell-type-specific subcellular structures and their boundaries (Fig. 1d). These informative representations will be used in the segmentation task as part of the multimodal input (Fig. 1e).

In the context of cell segmentation, traditional methods typically rely on pixel-level segmentation on images which, however, struggles to adequately represent the intricate relationships between adjacent transcripts and fails to discern cell assignments for transcripts at sub-pixel granularity. In contrast, to address these limitations, we have innovatively approached it as an edge prediction task, which significantly enriches the representations of gene relationships, in addition to the transferred physical colocalization relationships. This involves the classification of connections between nodes (transcripts or proteins) into two categories: those originating from the same cell and those originating from different cells (Fig. 1f).

Recognizing the pivotal role transcript colocalization in cell boundaries detection, we have devised a novel strategy, where we leverage the learned cell-type-aware node representations obtained from GNN and transfer them as edge embeddings for the edge prediction task (Fig. 1e, Supplementary Fig. 6). These embeddings are augmented with image-aware embeddings obtained from convolutional neural networks (CNN) and distance-aware embeddings derived from learnable radial basis function (RBF) kernels, respectively (Fig. 1e, Supplementary Fig. 6).

Additionally, multi-modal input with cell-type-specific node embeddings and RBF distance kernels provides flexibility for segmentation masks to adapt to varying sizes across different cell types, mirroring real-world scenarios (Supplementary Fig. 7). Following the predictions of edge labels, we apply community detection techniques such as Leiden clustering to accurately identify individual cells within the tissue slice (Fig. 1f) (Methods). This comprehensive process ultimately enables the precise segmentation of individual cells, which can be conveniently annotated using transcript-level cell annotations, streamlining downstream analysis. In the Bering model, the combination of node representations and distance kernels provides substantial knowledge for accurate image-free segmentation, particularly in sparsely populated tissues like the cortex.

Conversely, densely packed tissues such as tumors and the ileum can benefit from the Bering model that incorporates image-aware embeddings, improving segmentation performance, and thus, generating more accurate mapping of gene expression in space.

### Validating Bering performance of background noise and cell type prediction

Background noises pose a substantial challenge in some spatial technologies as they lack distinct boundaries from real signals, as exemplified in MERFISH and STARmap^3,20^. Bering addresses this issue by leveraging its GNN model to predict both background noises and real signals with cell-type annotations. To assess its performance, we conducted benchmarking experiments, comparing Bering with other methods for noise prediction and transcript-level cell annotation. In the case of mouse cortex MERFISH data, the original unsegmented transcripts accounted for more than half of the total transcripts, encompassing both noise and unsegmented real signals due to conservative segmentation in the original paper^3^. Baysor failed to accurately capture the real signals, whereas Bering demonstrated a more cautious approach in real signal prediction, resulting in more precise background noise prediction (Fig. 2a, Supplementary Fig. 8). In this case, Bering exhibited a notable improvement of up to 50% in noise prediction accuracy compared to other methods (Fig. 2b). Image-based segmentation methods such as Watershed and Cellpose often heavily rely on available staining images, particularly DAPI staining, which tends to yield conservative segmentation results (Fig. 2b, Supplementary Fig. 8b). Importantly, Bering consistently showcased superior performance in noise prediction across various technologies, including and MERFISH and pciSeq (Fig. 2b).

**Figure 2.**
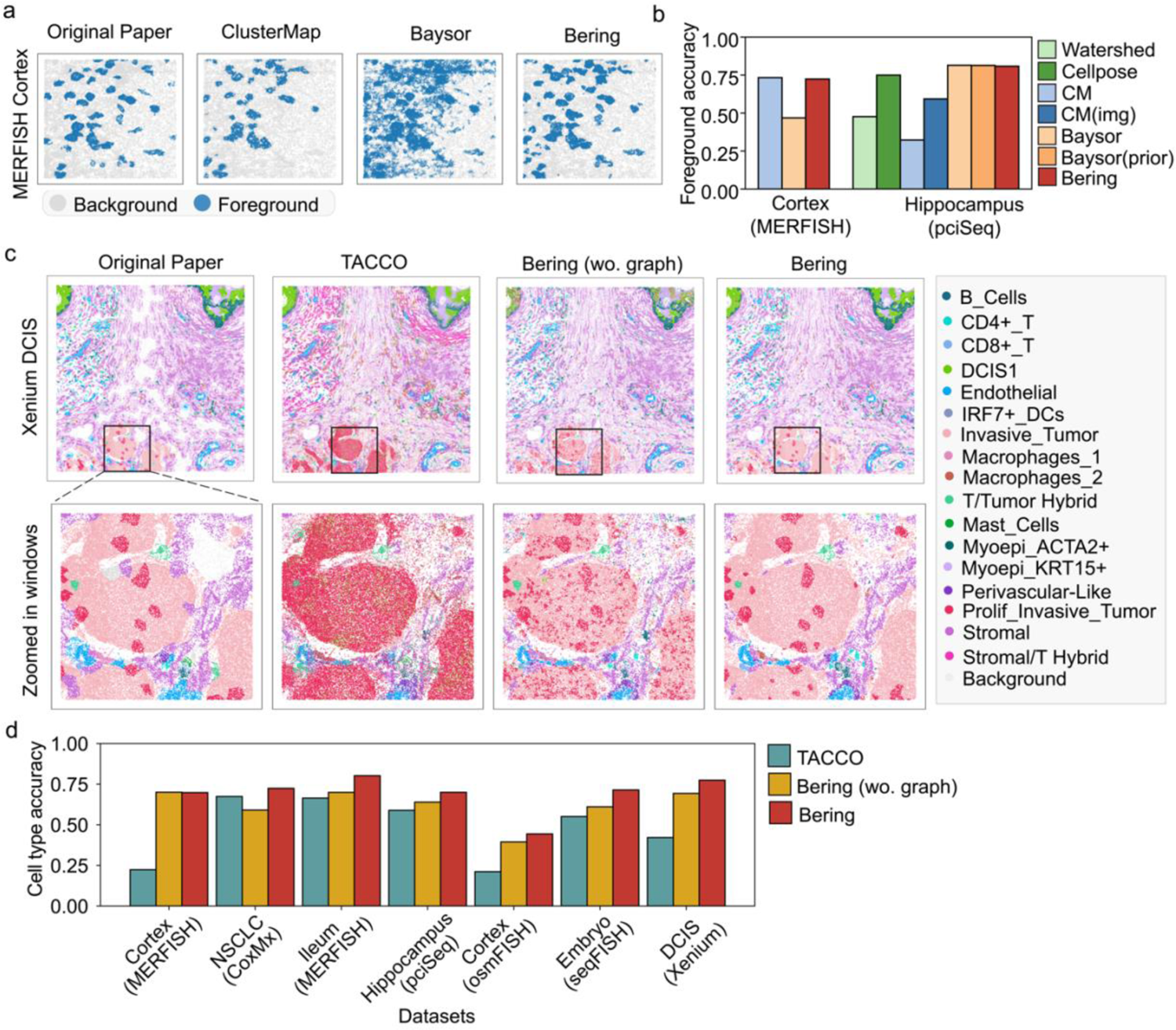
Performance of noise and cell type predictions on transcripts. (a) Background noise prediction in the MERFISH cortex data using different segmentation methods. The background noise annotated in the original paper of the data was shown on the left. (b) Bering demonstrates superior performance in predicting noise and real signals compared to other segmentation methods across datasets. CM: ClusterMap; CM(img): ClusterMap with DAPI image input. (c) Cell type prediction in the 10x Xenium data of Ductal Carcinoma In Situ (DCIS) using different transcript-level annotation methods, including TACCO and Bering with and without graph models (top). The zoomed-in visualization of a particular section of the tissue is presented below. (d) The accuracies of cell type prediction by TACCO and Bering, with and without graph models, were evaluated across various datasets.

Furthermore, we conducted a benchmark comparison of transcript-level cell type annotation using the state-of-the-art approach TACCO^21^. In the case of ductal carcinoma in situ (DCIS) Xenium data^9^, Bering’s predictions accurately identified cell labels and preserved more detailed tumor microenvironment components compared to TACCO (Fig. 2c, Supplementary Fig. 9).

Specifically, Bering successfully distinguished proliferative invasive tumor cells from other tumor cells in the niche, whereas TACCO failed to differentiate between these two types of tumor cells (Fig. 2c). Additionally, Bering captured more comprehensive immune cell distributions within the tumor microenvironment in non-small cell lung cancer (NSCLC) (Supplementary Fig. 9c). Importantly, Bering with graph models demonstrated fewer sporadic predictions and more consistent cell predictions compared to Bering without graph models, highlighting the advantages of information sharing in the neighborhood facilitated by graph models, which aligns with our initial hypothesis during model construction (Fig. 2c, Supplementary Fig. 9a). Implemented across diverse technologies and tissues, Bering consistently achieved higher accuracy in cell type prediction, with improvements ranging from approximately 10% to 50% compared to TACCO (Fig. 2d).

### Validating Bering performance on cell segmentation

The Bering model for cell segmentation incorporates various components, including graph models, RBF kernels, and image embeddings learned from CNNs. To assess the contribution of each module and understand the model’s capabilities, we conducted ablation studies and evaluated the performance of cell segmentation using quantitative metrics such as adjusted mutual information (AMI), the fraction of assigned molecules, and the number of segmented cells. The results revealed that the inclusion of either image embeddings or RBF distance kernels led to significant improvements in segmentation accuracies (Supplementary Fig. 10a) and sensitivities (Supplementary Fig. 10b). While image-free segmentation performed well in the CosMx NSCLC data, the addition of image embeddings increased the number of segmented cells by approximately 10% (Supplementary Fig. 10c). Consequently, for most datasets, we implemented Bering with graph models and RBF kernels, and incorporated image embeddings if cell staining images were available.

Prior to conducting comprehensive benchmark studies, we performed a hyperparameter search for both Bering and the benchmark methods (Supplementary Fig. 11-14). In Bering, we thoroughly compared hyperparameters such as the number of GNN layers, number of training cells, and structures of RBF distance kernels to determine the optimal settings (Supplementary Fig. 11). In the cell segmentation process of Bering, unsupervised clustering is involved, and the hyperparameter of clustering resolution can be set manually. It was observed that stable cell segmentation results were achieved when the edge prediction accuracy was high (Supplementary Fig. 12). This implies that stable segmentation can be obtained by focusing on improving the accuracy of edge prediction, rather than purely adjusting the clustering resolution hyperparameter. Additionally, we searched hyperparameters for the benchmark methods to achieve the best segmentation performance for benchmark studies (Supplementary Fig. 12-13).

We then implemented the benchmark methods on the NSCLC CosMx data and observed that Bering accurately preserved cell boundaries compared to other methods. In contrast, Watershed and Cellpose exhibited a relatively conservative segmentation approach, while ClusterMap and Baysor predicted a certain number of cells with abnormal sizes (Fig. 3a). Similar observations were made in other tissues, including ileum, cortex, and DCIS (Supplementary Fig. 15).

**Figure 3.**
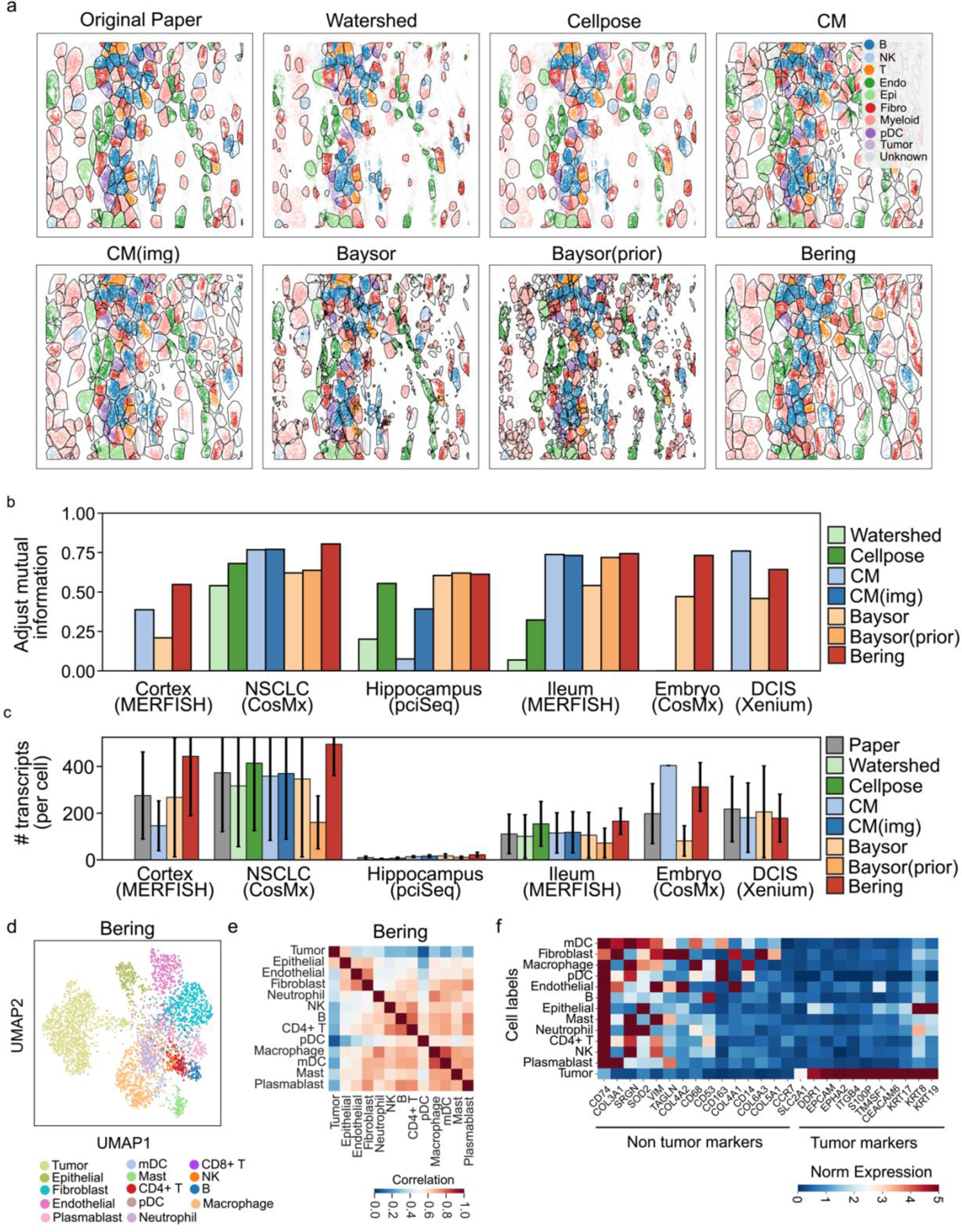
Performance of cell segmentation. (a) Zoomed-in sections of CosMx NSCLC data illustrate the cell segmentation results obtained using various segmentation approaches. Different cell types are depicted in distinct colors, while background noises are visualized as gray dots. Cell boundaries are depicted using hulls. The segmentation result from the original paper is displayed in the top-left corner. CM: ClusterMap. CM (img): ClusterMap with DAPI image input. (b-c) Quantitative metrics, such as adjusted mutual information (AMI) (b) and the number of transcripts per cell (c), are employed to benchmark the segmentation results across diverse datasets. The error bars represent the standard deviations of transcripts per cell. Image-dependent methods were excluded from the benchmark if processed nuclei staining images were unavailable. (d) The UMAP shows single cells and labels generated from Bering segmentation results. (e) Spearman correlation calculated for gene expression levels in cell labels shown in (d). Representative cell markers, obtained through differential expression analysis from the original single-cell data in the paper, were utilized for correlation measurement (See Methods). (f) Expression levels of tumor and non-tumor genes across cell types in (d). Additional single-cell analysis and correlation matrices from other segmentation results can be found in Fig. S17 for comparison.

Quantitative measurements also indicated the superior segmentation performance of Bering in terms of prediction accuracies across different datasets and technologies (Fig. 3b). Moreover, Bering consistently detected a higher number of transcripts in individual cells with a relatively lower standard deviation, indicating a higher signal detection capacity and stable segmentation sizes (Fig. 3c). This observation is supported by the measurement of segmented cell areas and the fraction of assigned molecules, where Bering segmented cells with larger average sizes (up to 40% compared to the original paper) and harvested more transcripts during the segmentation process compared to other methods (Supplementary Fig. 16).

To gain insights into the quality of single cells derived from different segmentation methods, we conducted benchmark comparisons at the single-cell level using the NSCLC CosMx data, where cell labels obtained from model predictions or label transfers were displayed within reduced dimensions. (Fig. 3d, Supplementary Fig. 17a, Methods). We measured the correlations between cell types. Remarkably, we observed that Bering exhibited low correlations between tumor and non-tumor cells, and closely mirrored the correlation patterns observed in the original paper (Fig. 3e, Supplementary Fig. 17b). In contrast, other methods demonstrated strong cross-correlations between tumor and non-tumor cells, alongside diminished correlation within non-tumor cells (Supplementary Fig. 17b). These findings suggest that Bering’s segmentation results have cleaner captured signals and less contamination. Furthermore, the expression of marker genes clearly indicates the separation of tumor and non-tumor cells (Fig. 3f).

### Bering’s applications in versatile spatial technologies and 3D thick tissues

A diverse range of technologies now exists for generating single-cell spatial transcriptomics data, offering distinct data qualities and gene throughput capabilities. For instance, osmFISH enables the capture of 35 genes, while seqFISH+ has scaled throughput to accommodate up to 10,000 genes (Supplementary Fig. 1, Table S1). However, the application of these technologies across different tissues presents significant challenges in terms of cell segmentation^2^. To showcase the effectiveness of Bering, we have applied the model to various tissues, including densely packed tissues like the ileum and embryo (Fig. 4a-h). Our model generates precise cell boundaries with corresponding cell annotations, providing a convenient resource for downstream analysis. A holy grail of spatial transcriptomics is to generate spatially resolved gene expression in 3D tissues and organs, thus, we applied Bering to the latest 100-µm thick-tissue MERFISH cortex tissue^22^ for 3D segmentation (Fig. 4i). We segmented 397 cells and harvested 530,912 transcripts, 9.3% higher than the original paper. In summary, Bering successfully segments cell boundaries and accurately predicts their corresponding cell types, thereby demonstrating its efficacy in simultaneous cell segmentation and annotation across diverse datasets and technologies, in both 2D and 3D settings.

**Figure 4.**
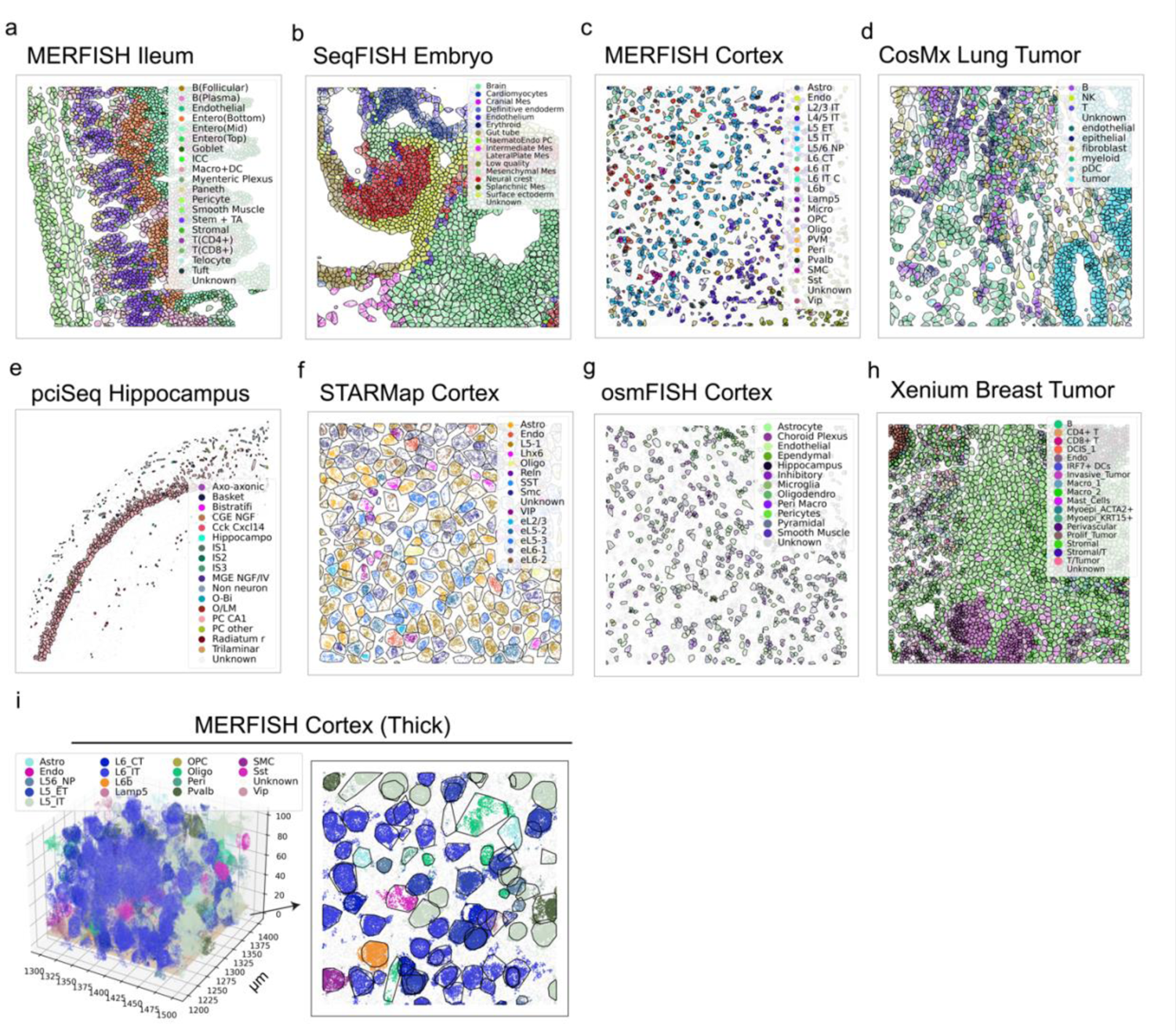
Bering applications across technologies and tissues. (a-h) Bering was applied to various single-cell image-based spatial datasets, with predicted cell types and boundaries depicted in different colors and hulls in zoomed-in regions. Predicted background noises were colored in light gray. (i) Bering was applied to thick-tissue MERFISH mouse cortex dataset, resulting in the prediction of diverse cell types (left) and the successful segmentation of individual cells. A cross-section at 10 µm (±5 µm) was magnified and presented on the right, highlighting the segmentation outcome for a specific plane.

### Generalized Bering model using transfer learning and self-distillation

Bering, as a deep learning approach, provides the distinct advantage of cross-dataset portability. For example, we successfully applied a pre-trained model developed from one slice of the mouse cortex to another slice, yielding highly satisfactory segmentation performance. This led to unambiguous cell type annotations (Supplementary Fig. 18a) while achieving comparable performance to the fine-tuned model (Supplementary Fig. 18b).

However, the portability of the Bering pre-trained model can be significantly hindered by batch effects across datasets, where the throughput of genes, which serve as features in the node classification task, can vary dramatically. This presents substantial challenges when applying pre-trained models to new data. To overcome this obstacle, we employed transfer learning techniques and employed the self-distillation method to enhance prediction results on the new data (see Methods). In the specific case of the cortex MERFISH data, we acquired a pre-trained Bering model from Zhang et al.^3^ and utilized it to predict cells and annotations in the new cortex VISp data from Biancalani et al.^23^. Less than 20% of molecules were assigned cell labels initially by the pre-trained model (Fig. 5a-c). To better capture the latent representation of the new data, we improved the pre-trained model through two rounds of self-distillation, leveraging the coarse prediction labels in the new data (see Methods). As a result, a larger number of transcripts were successfully labeled and segmented, with over 80% of transcripts assigned labels and more than 2,000 cells segmented (Fig. 5c). Notably, the different layers of neurons (L2-L6) were accurately predicted, with intermittent distributions of interneurons and supporting cells (Fig. 5a, b). Furthermore, the predicted single cells from various cell types exhibited distinct distributions on the UMAP, highlighting a more pronounced separation between cell types compared to the predictions prior to self-distillation.

**Figure 5:**
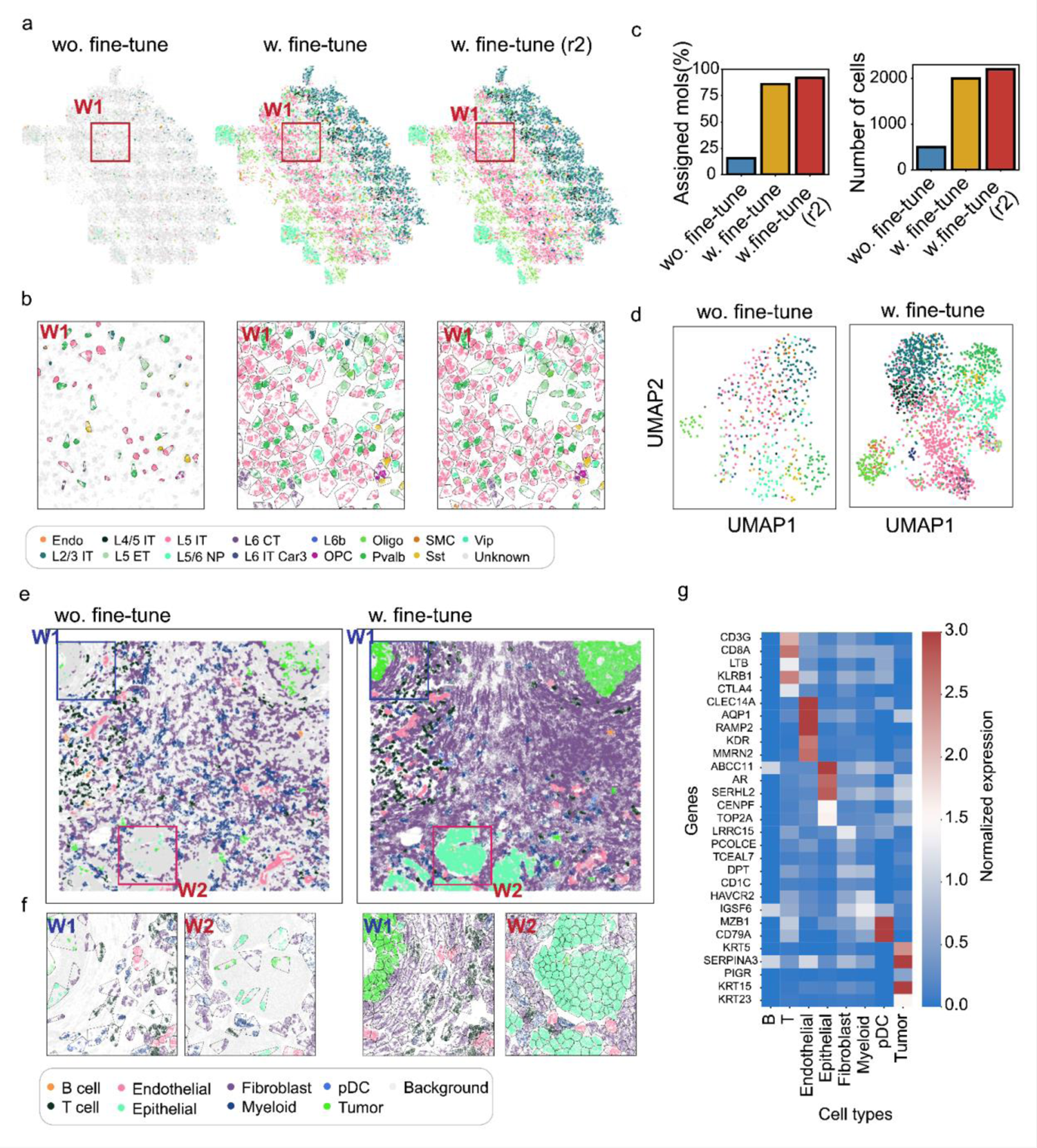
Generalizability of the Bering model using transfer learning and self-distillation. (a-d) Transfer learning of the Bering pre-trained model on a new mouse cortex MERFISH dataset. (a) Application of a pre-trained model from mouse cortex MERFISH data (Zhang et al.) to new mouse cortex data (Biancalani et al.), with and without fine-tuning. Fine-tuning labels were derived from the prediction results of the pre-trained model, shown in the leftmost figure. Two rounds of fine-tuning were conducted through distillation of the prediction results (see Methods). A specific region is highlighted for further investigation. (b) Enlarged view of the highlighted region in (a). Cell boundaries are depicted by hulls. (c) Quantitative metrics depicting the increasing percentages of assigned molecules (left) and the number of segmented cells (right). (d) UMAPs displaying the distributions of cells from prediction results, with and without fine-tuning. (e-g) Similarly, transfer learning of Bering on tumor spatial transcriptomics data. (e) Application of a pre-trained model from CosMx NSCLC to Xenium DCIS data, resulting in prediction results without and with fine-tuning, respectively. Two specific regions are highlighted for further investigation. (f) Enlarged views of the highlighted regions in (e) showing predicted cell types and cell boundaries in the results with and without fine-tuning. (g) The heatmap shows the expression levels of marker genes in the segmented cells from the tumor microenvironment.

We further applied this strategy to tumor datasets, where the pre-trained model was derived from NSCLC CoxMx data comprising 980 genes, and the validation data was obtained from the DCIS dataset, consisting of 313 genes. Without any fine-tuning, approximately 40% of transcripts in DCIS were successfully segmented and assigned cell labels (Supplementary Fig. 18c). However, the resulting transcript distribution landscape on the tumor slice lacked sufficient details for individual cells (Fig. 5e,f). Through the process of self-distillation, we achieved significant improvements. More than 80% of transcripts were labeled, and over 3,000 cells were successfully segmented, which is approximately three times more compared to the results before self-distillation (Supplementary Fig. 18c). This enhancement allowed us to reveal finer details of tumor niches, such as the colocalization of immune cells and tumor cells, as well as the precise boundaries of tumor regions (Fig. 5e,f). Notably, the expression patterns of marker genes demonstrated specific expression across different cell types, confirming the accuracy of our cell segmentation and annotation.

## Discussion

Cell segmentation remains a challenging task today for several reasons. One of the primary difficulties is that some tissues have densely packed cells with unclear boundaries, making it difficult to make accurate segmentation. For instance, some cells in tumor tissues and ileum^17^ have almost no gaps between them, presenting a very different scenario than the cortex, where tissues have a sparse distribution of cells. Additionally, the limitations of capture rates and RNA diffusion in some technologies, such as pciSeq and MERFISH^6^, result in sparse and noisy spots. Cell staining imaging, including DAPI, poly-A, and membrane staining, reveals imbalanced image signals among different cell types and incomplete capturing of entire cells due to limitations in scanning depth. While traditional methods have struggled to achieve accurate cell segmentation masks, the molecular information generated by spatial technologies has opened a new avenue to address this challenge.

The field of subcellular transcriptomics has gained popularity as a result of the rapid advancement of spatial omics technologies^11,14,15,19,24^. These cutting-edge technologies have enabled researchers to obtain more detailed information about cellular processes than ever before. For example, recording the transmission of neurotransmitters has been difficult in the past, but with high-resolution technology such as Ex-seq^7^, it is now possible to study neuron interactions within the synapse. Over the last few years, high-throughput sequencing-based spatial technologies, such as Slide-tag and Stereo-Seq have greatly enhanced the spatial resolution to near-single-cell or subcellular levels^25^. Additionally, the size of features in image-based spatial transcriptomics technologies has increased from 30 to 10,000^4^, making it increasingly feasible to use deep learning models. However, the primary obstacle to implementing such models remains the shortage of labels for specific tasks. Consequently, previous methods for analyzing subcellular data, such as Bento and FISHFactor, have primarily relied on statistical models for unsupervised learning^15,19^. However, cell segmentation, which is relatively easier to obtain a large number of labels using traditional methods, presents the possibility of exploring the application of deep learning models in subcellular resolutions. Benchmark results in our study demonstrate the superior performance of deep learning models compared to traditional methods by inferring the complicated underlying relationships of gene colocalizations.

In our paper, we explore the possibility of detecting cell borders using subcellular transcript distributions and demonstrate the successful application of the graph model. Although Bering gained good performance in cell annotation and segmentation, we expect Bering can be further improved in future studies. First, graphs in the model were built using *k*-nearest neighbors, which mainly consider the relationships of transcript location within a local region. However, cells are usually sphere shapes in their 3D tissue environments and the relationships of transcripts on the membrane of two sides of the cell may not be efficiently captured by the graph. Second, the model requires labels for training purposes. If the labels come from Watershed, which may over-rely on the nuclei position and give conservative cell masks, it may also lead to conservative prediction in the Bering model. Third, the speed of training for the image-based encoder is relatively slow compared to image-free segmentation, where we only use the gene-colocalization information as the input. Technical performance can still be improved for better and faster training and prediction.

Obtaining pre-trained labels can be a difficult and labor-intensive task. Consequently, it would be advantageous if we could obtain pre-trained models for specific tissues based on large-scale data. The cortex^20^, with its accessible subcellular transcriptomics data, is an example of such a tissue. We have demonstrated that a pre-trained model can be readily applied to a new dataset or a new technology of the same tissue and enhanced through fine-tuning the data itself. However, the main hurdle in obtaining practical pre-trained models for various technologies is the variability in the features being measured and the lack of a large amount of data. For instance, the overlap of measured genes between the MERFISH cortex data and the osmFISH cortex data is limited to less than 15. This presents a significant challenge for transfer learning in such cases. The selection of genes to be probed and measured is heavily influenced by authors’ interests, which can result in significant heterogeneity across datasets. The varying choices and depth of features create challenges for portable models. Nevertheless, as more genes are probed in new versions of these technologies and more datasets from various tissues are generated, this issue will become less problematic in the future.

In summary, we tackled the cell segmentation task for spatial data using a graph model and transfer learning, incorporating multi-modality information during training. We demonstrated the model’s portability across various datasets and technologies. Nonetheless, there are still some challenges that we hope to address with future technological advancements.

## Methods

### Bering model framework

The Bering model, illustrated in Figure 1 and Supplementary Fig. 7, consists of two main components. The first component involves node classification to distinguish between noise and real signals, with predictions made for cell types. In the second component, cell segmentations are performed, which entails predicting edges. Initially, we construct a gene colocalization graph, which serves as the input for a graph neural network used in the node classification task. The node representations, along with auxiliary edge embeddings derived from image staining and distance kernels, are employed as edge embeddings. These combined edge embeddings are utilized to predict edge labels, wherein intercellular and intracellular edges are binarized as negative and positive labels, respectively. The final edge predictions are then utilized to build a molecular connection graph, where a community detection algorithm is applied to achieve cell segmentation. Finally, the outcomes of both node classification and cell segmentation tasks are merged to obtain annotated single-cell data.

### Gene colocalization graph

In this model, molecules, such as transcripts, are depicted as nodes on the slice, and we utilize their 2-dimension (2D) or 3-dimension (3D) spatial coordinates to construct *k*-nearest neighbor graphs (KNN) to capture gene colocalization information. By default, we consider the 20 nearest neighbors. Edges of the graph depict equal-weight neighborhood relationships between nodes.

The graph is described as below:

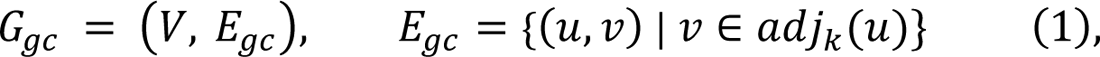

where *V* represents the node set, *E*_*gc*_ represents the edge set and *adj*_*k*_(*u*) represents the k-nearest neighboring nodes of node *u* in Euclidean space.

### Node features

To capture gene colocalization information in 2D or 3D spatial coordinates more effectively, we utilize neighborhood gene components (NGCs) as node features. NGCs consist of gene components within the k-nearest neighborhoods, resulting in sparse count matrices where genes serve as features. This enables us to incorporate spatial relationships and uncover insights about gene colocalization patterns. Below is the definition of NGC:

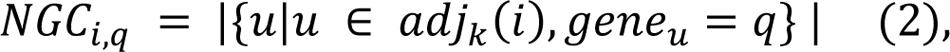

where *i* represents molecule *i* in the node set *V* and *q* denotes gene *q* in the gene set. The matrix value *NGC*_*i,q*_ indicates the count of detected genes *q* in the neighborhood of node *i*. The total number of genes in the dataset is defined as *N*_*genes*_, and each gene *u* corresponds to a column in the NGC matrix. The matrix values indicate the count of detected genes *u* in the NGC. *N*_*genes*_ can vary across technologies and datasets. For instance, osmFISH has been reported to detect 35 genes^26^, whereas SeqFISH+ has the capacity to detect up to 10,000 genes in a single experiment^4^.

### Graph convolutional networks

Assuming that spatially proximal nodes exhibit similar node embeddings, our hypothesis aligns with graph neural networks. These networks propose that the node representation in a graph should consider not only its own features but also the characteristics of its neighboring nodes. In our model, we employed the Graph Convolutional Network (GCN) to analyze the gene colocalization graph. The GCN layer is defined as follows:

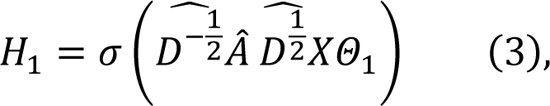

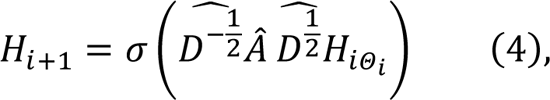

where *H* represents the matrix of node representations ℎ_*u*_, and *X* represents the NGC of node features *x*_*u*_. σ(·) denotes the activation function (ReLU in our case). Â is the graph adjacency matrix augmented with self-loops. Ď is the graph degree matrix, and Θ is a matrix of trainable parameters.

### Fully connected neural networks

After obtaining the node representation from GCNs, Fully Connected Networks (FCNs) are employed. The FCN is defined as

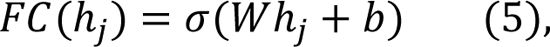

Here ℎ_*j*_ denotes the representation of node *j*. The weight matrix *W* and bias term *b* are learned for each layer. σ(·) denotes the activation function (ReLU in our case). Multiple FCN layers are stacked to get the final prediction. Similar networks are employed for edge predictions in the segmentation task as well, which will be mentioned below.

### Node classification

Node label prediction is accomplished by leveraging the node representations acquired through GCNs and FCNs. In the training phase, the objective function is determined by computing the cross-entropy loss between the ground truth labels and the predicted labels.

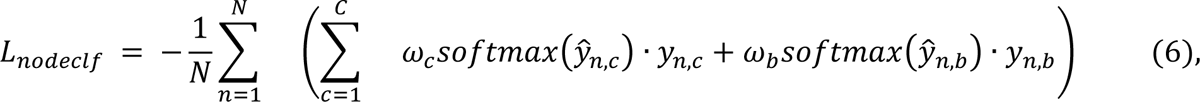

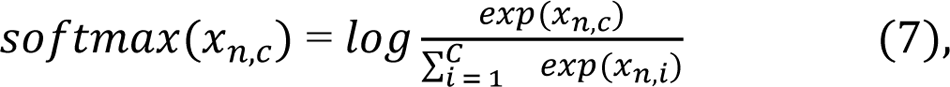

where the loss of node classification is denoted as *L*_nodeclf_. ŷ ∈ *R*^|*V*|×(*c*+1^) is the output from FCNs and *y* is the ground truth labels. *C* is the number of classes, and ω is the weight. ω_*c*_ and ω_*b*_ represent the weight of real signal nodes with various cell types and background noises, respectively. These weights can be adjusted by users according to the background noise prevalence to effectively identify real signals and noises.

### Transfer node representation for segmentation

NGCs are employed to learn the node representations of molecules, primarily for the node classification task. However, our ablation study revealed that this representation proves advantageous for edge classification in the cell segmentation task as well. Consequently, we transfer the node representations acquired from NGCs and concatenate the representation matrix of two end nodes of an edge, forming a new matrix that becomes a part of the edge representation. Throughout this process, the parameters of GCNs and FCNs learned for the node classification task remain frozen.

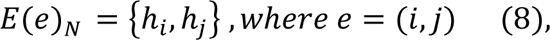

where *E*(*e*)_*N*_ represents the edge embedding of edge *e* by concatenating node representation ℎ_*i*_ and ℎ_*j*_ of node *i* and *j* from the layer *l* of the node classification model. By default, we select the output of the first fully connected layer after GCNs as the node representation.

### Distance kernels

The RBF distance kernel is a kernel function that measures the similarity between two vectors based on the distance. Since the spatial distance between two nodes is highly correlated with their intercellular relationship, we utilize distance kernels as a part of the edge embeddings to effectively learn the appropriate cell sizes.

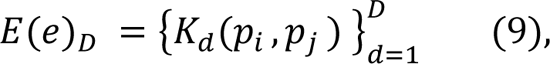

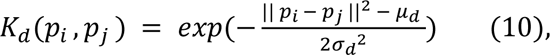

where *E*(*e*)_*D*_ represents the edge embedding derived from distance kernels *K*_*d*_. Total number of kernels is denoted as *D*. *p*_*i*_ and *p*_*i*_ represents coordinates of nodes *i* and *j*. μ_*d*_ and σ_*d*_ are the mean and standard deviation of each kernel, which can be learnable.

### Image representation

Cell boundaries are highly indistinct without image staining in densely packed tissue, such as tumor and ileum. Latent image representations prove beneficial in these cases. Consequently, the image representations serve as an additional edge representation in such scenarios. In this model, CNNs followed by spatial pyramid pooling (SPP) are applied to learn the embeddings of input images of different sizes. FCNs are further utilized to learn the edge representation using the output of SPP.

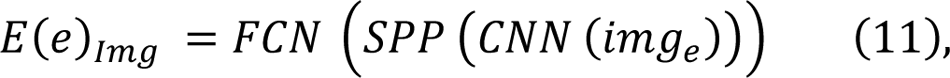

where *E*(*e*)_*img*_ represents the image representation in the edge embedding, and *img*_*e*_ is the image of edge *e*. The image of an edge is defined as the rectangular region it covers, with the edge itself forming the diagonal line. While the size of these images falls within a specific range, the exact dimensions can vary significantly (such as 7 x 12 or 14 x 19 pixels), depending on the length of the edge. To optimize computational resources, we categorize edge sizes into distinct groups (such as 5 x 10 or 15 x 20 pixels). The actual images are then adjusted by expanding or cropping to fit within the designated bin size, which serves as the training input.

### Edge representation

The edge representation serves as the input for the edge classification and the segmentation task, comprising three aforementioned key components, including node representation, distance kernels and image representation.

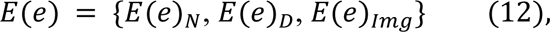

These three components play a crucial role in efficiently learning the underlying knowledge of gene colocalization relationships, cell sizes, and image-informed cell boundaries, respectively. Moreover, their combination holds the potential to learn cell type-specific cell sizes. Node representations and distance kernels prove to be adequate for sparsely populated tissues with clear cell boundaries, such as the cortex. However, in densely packed tissues like tumors, ileum, and liver, where cell boundaries are often challenging to discern, image representation becomes a vital component. The incorporation of image channels, such as DAPI staining, provides valuable information about cell boundaries in these densely packed tissues.

### Edge classification

The edge classification task serves as input for the cell segmentation community detection algorithm. It is formulated as a binary classification problem, training the fully connected neural network to discern intracellular and intercellular molecular colocalization. Predicted edge labels are obtained using the sigmoid function applied to the neural network output. Binary cross-entropy is employed as the objective function.

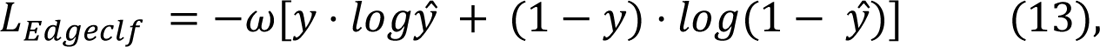

where *L*_*Edgeclf*_ represents the loss of the edge classification model. ω is the weight; *x* and *y* are the probability of predicted binary labels and ground truth labels, respectively.

### Molecular connectivity graph

After the edge prediction, we create the molecular connectivity graph using predicted edge labels *e*_*i*,*j*_. Positive labels (*e*_*i*,*j*_ = 1) indicate a connection between two nodes, where they belong to the same cell according to the model. Negative labels (*e*_*i*,*j*_ = 0) imply no connection between nodes, indicating they belong to separate cells.

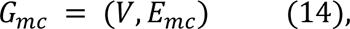

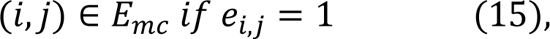

Due to the large number of nodes and edges involved (potentially in the millions and billions, respectively), it is infeasible to predict labels for all pairwise edges. Currently, technology typically covers fewer than 300 nodes in a single cell, suggesting that positive edges for a node primarily come from its 300 closest neighbors. Consequently, we only predict edge labels for nearest neighbors.

### Cell segmentation with community detection algorithms

Once the molecular connectivity graph is constructed, we apply the community detection algorithms, such as Louvain and Leiden, to identify clusters from the graph. These clusters correspond to individual cells in the tissue.

### Single-cell annotation

After obtaining node classes and cell boundaries, we utilize both outcomes to generate single-cell data with annotations. We select cells that meet specific criteria, including (1) a minimum number of total transcripts per cell, (2) a minimum number of transcripts for the dominant cell type per cell, and (3) a minimum ratio of transcripts of the dominant cell type per cell. We introduce a threshold for the transcripts of the dominant cell type as we posit that transcripts within a segmented cell should belong to the same cell type. If transcripts within a segmented cell are annotated as many different cell types, we lack confidence in the accuracy of the cell annotation. By default, these thresholds are set as 50, 30, and 0.6, respectively. Segmented cells that satisfy the criteria are annotated with the dominant cell class of transcripts within each cell.

### Model implementation

The GCNs are implemented using PyTorch Geometric^27^, with batch normalization and dropout layers (dropout rate = 0.2) applied to each graph convolutional layer during training. The FCNs and CNNs are implemented using PyTorch, with batch normalization incorporated into each FCN layer. To handle the heterogeneous shapes of input images, spatial pyramid pooling is employed. This is achieved through the PyTorch AdaptiveMaxPool2d function, utilizing three pooling sizes (4, 2, and 1) to ensure consistent sizes of output embeddings. The resulting embeddings from the spatial pyramid pooling layers are passed through two layers of FCNs to generate the image representation, which becomes a part of the edge representation. The RBF distance kernel, referred to as the GaussianSmearing function, is implemented using the TorchDrug package^28^. The parameters μ and σ in the distance kernels are learnable during the training phase.

In both the node classification and edge prediction tasks, a learning rate of 1 x 10^-3^ and weight decay of 5 x 10^-4^ are set. Early stopping can be triggered when the loss fails to decrease.

### Generalized model using transfer learning

Fine-tuning is essential when working with new data that exhibits observable variations, as it helps enhance the model’s performance in both node classification and edge prediction. By retraining pre-trained Bering models using new data from the same tissue, the model parameters are updated using the training data and labels from the new dataset. This process ensures that the model adapts to the specific characteristics and patterns present in the new dataset, thereby improving its overall performance.

### Analysis of Neighborhood Gene Component

To unravel the underlying information regarding gene colocalization, we construct atlases of neighborhood gene components (NGCs) using the CoxMx NSCLC dataset and the MERFISH cortex ileum dataset. In each cell, a random selection of two molecules is made, and their k-nearest neighbors are identified, resulting in the formation of two NGCs. These NGCs were derived from various cell types and compartments, including nuclei, cytoplasm, and membrane. The information about cell type and compartment is derived from the original papers. The NGCs from different cell types and compartments are then concatenated to form a matrix, similar to single-cell matrices where genes serve as features. The distinction lies in the fact that NGCs utilize neighborhoods as objects within the matrix, rather than individual cells. These NGC matrices are stored as Scanpy objects^29^, and an analysis pipeline resembling single-cell analysis is applied to the data. By employing UMAPs and Leiden clusters, we obtain reduced dimensions and clustering patterns from these matrices. For further details, please refer to the single-cell analysis section.

### Ablation study

First, we employed baseline fully connected neural network (FCN) models and graph convolutional network (GCN) models with identical layer dimensions for the node classification task. We evaluated the classification accuracies across multiple datasets, as depicted in Figure 2c, d and Supplementary Fig. 9. Furthermore, we conducted eight ablation studies on the NSCLC CosMx data to assess the impact of different model components, such as GCNs, RBF distance kernels, and image embeddings. All other model hyperparameters remained constant. The ablation studies involved measuring various segmentation metrics, including adjusted mutual information (AMI), fractions of assigned molecules, and the number of detected cells.

### Benchmark metrics

#### Node classification task

In this project, we employed multiple metrics to evaluate the performance of the model from various angles. For the node classification task, we focused on two aspects for comparison: background noise prediction and cell type prediction. In in-situ hybridization methods like MERFISH, significant noise is observed due to RNA diffusion during the staining rounds. To assess the effectiveness of background noise classification, we utilized accuracy as the evaluation metric. We obtained the ground truth by extracting foreground real signals and background noise, and then calculated the accuracy using the predicted labels generated by the model.

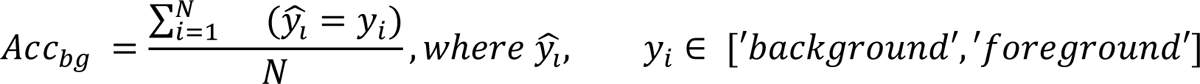

*N* is the total number of molecules, and ^ŷ^_*i*_ and *y*_*i*_ are predicted labels and ground truths of real signals and background noises for the molecule *i*. Apart from background accuracy, we also evaluated the performance of cell type classification. The accuracy of cell type predictions is used for the comparison,

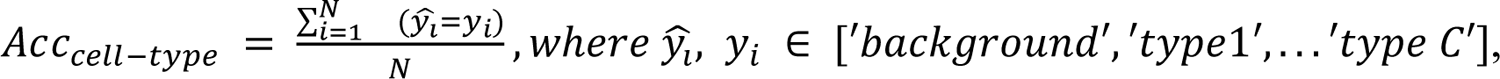

where ^ŷ^_*i*_ and *y*_*i*_ are *predicted labels and ground truth* of cell types for the molecule *i*.

#### Cell segmentation task

The cell segmentation task can be seen as an unsupervised clustering problem, where the similarity between our predicted cells and the cells in the ground truth needs to be assessed. To measure this similarity, we employed a widely used metric known as adjusted mutual information (AMI). AMI served as the quantification metric in evaluating the performance of our cell segmentation method.

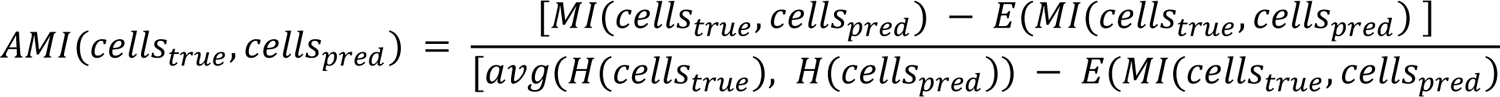

where *cells*_true_, *cells*_pred_ represent cell ids from ground truth and predictions. *H*(·) represents the entropy of a vector, and *MI*(·) represents mutual information. AMI considers the fact that mutual information is generally higher for two clusterings with a large number of clusters. In our case, the number of cells in a dataset is usually large and AMI could be a more appropriate option.

#### Fraction of assigned molecules

The fraction of assigned molecules is utilized as a metric to compare the effectiveness of segmentation. We hypothesize that more conservative methods that solely rely on nuclei staining images may yield lower performance in this metric. It is defined as below.

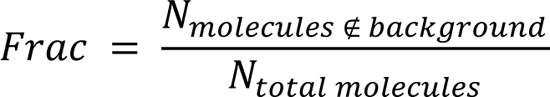

#### Number of cells and cell areas

The number of cells is utilized as an indicator to assess the capability of identifying individual cells. Nevertheless, it is important to note that a higher number of cells does not necessarily indicate superior segmentation performance. Certain methods may employ an aggressive approach that results in numerous cells with small areas. Therefore, we also evaluated the sizes of the segmented cells to determine whether they exhibit appropriate shapes.

#### Correlation of single cell expressions

To assess the correlation between clusters, we initially identified representative cell markers by conducting differential expression analysis within the single-cell clusters from the original paper^8^. Subsequently, we calculated the Spearman correlation by averaging the gene expression values across clusters. This allowed us to evaluate the degree of correlation between the clusters.

### Benchmark methods

#### TACCO classification

TACCO, an optimal transport method, enables the transfer of annotations from single-cell data to spatial data. In our benchmark studies, we utilized the benchmark datasets with annotations obtained from the original papers to create the reference single-cell data. The cell type annotations for molecules were generated by projecting the single-cell labels using the “annotate_single_molecules” function.

#### Watershed segmentation

The watershed method, implemented using the spateo package^30^, was utilized for segmentation based on nuclei staining images with some modifications. Firstly, the masks of nuclei were identified using both global and local adaptive thresholding techniques. Subsequently, peak detections were performed based on the results obtained from the distance transform algorithm. The connected peaks were merged to form individual markers. Finally, the masks and markers were used as input for the watershed algorithm. Molecules were assigned to the nearest pixels, and cell IDs were determined based on the corresponding pixels.

#### Cellpose segmentation

Cellpose, a U-Net based deep convolutional neural network approach, was used as a benchmark method in the paper. We used the pretrained model ‘cyto’ and nuclei staining for segmentation.

#### ClusterMap segmentation

ClusterMap is a cell identification method that utilizes density peak clustering of spots in spatial data, with the option to incorporate staining images as an auxiliary input. In our benchmarking, we evaluated both modes of ClusterMap, namely with and without aligned nuclei images.

Specifically, we employed the 2D segmentation mode for this particular task. When available publicly, DAPI staining imaging was employed during the preprocessing stage. In cases of noisy data, such as MERFISH^3^ and pciSeq^6^, the noise ratio was estimated by referencing the percentage of unsegmented transcripts as indicated in the original paper; this parameter, termed “pct_filter,” was then configured during preprocessing. Furthermore, for these noisy datasets, the local noise rejection mechanism (“LOF”) was activated during the preprocessing phase.

#### Baysor segmentation

Baysor is a Bayesian model-based method designed for segmenting spots in spatial data. It has the capability to incorporate prior segmentation masks, such as those obtained from Watershed or other segmentation methods. In our benchmark study, we evaluated the performance of Baysor both with and without prior segmentation. For the tests with prior information, we performed Watershed segmentation on the nuclei staining images and utilized the resulting masks as input for Baysor.

#### Hyperparameter tuning

To ensure a fair benchmark comparison, we carefully fine-tuned the hyperparameters of Bering and other benchmark methods, selecting the best parameter combinations for the final evaluation. In the case of the Watershed algorithm, we considered three important parameters: the minimal distance for peak detection, the kernel size for morphological open and close operations, and the block size for adaptive thresholding. For Cellpose, we conducted benchmarking experiments to determine optimal diameters and flow thresholds. In Clustermap, we separately benchmarked two modes: one with auxiliary images and one without. In both modes, we evaluated three hyperparameters, including thresholds for cell numbers, estimated radius in the x-y plane, and sample interval in the DAPI image. Similarly, in Baysor, we evaluated its performance in two modes: one with a prior segmentation mask and one without. For Baysor, we first benchmarked hyperparameter selections, including scale, the standard deviation of scale, and the minimal number of molecules. Once we obtained the best hyperparameter combination, we applied it to the Baysor mode with a prior segmentation mask and further benchmarked the confidence of the prior segmentation. Lastly, for Bering, we benchmarked several hyperparameters, including the number of layers, the number of cells for pretraining, neighborhood sizes of graphs, and the numbers and size ranges of RBF distance kernels. The benchmark experiment across segmentation methods was conducted using the best hyperparameter selections we obtained.

#### Thick tissue analysis

The thick tissue MERFISH cortex dataset^22^ was harnessed to assess the 3D segmentation performance of Bering. The original cell boundaries and labels served as the pre-trained references during the model’s training phase. By employing Euclidean distances within the 3D context, we computed *k*-nearest neighbors and subsequently established colocalization graphs. The trained model’s prowess was then put to the test in a designated 200 x 200 x 100 µm region for comprehensive evaluation.

#### Subcellular pattern identification

Utilizing FISHFactor, we discerned distinct subcellular gene patterns within tumor cells from the NSCLC CosMx dataset. A cohort of 200 tumor cells was randomly selected, and three distinct factors were computed for all genes. The gene weights within these factors, along with spatial factor scores, were subsequently plotted to unveil subcellular gene patterns. Factor 2 and factor 3 were denoted as “nucleus-pattern” and “peripheral-pattern,” respectively. The genes with the highest weights within these two factors were identified, and their subcellular distributions were visualized. To portray these distributions (Fig. 1c), boundaries for cell nuclei and cell bodies were established through Cellpose segmentation of DAPI imaging and the convex hulls of all transcripts, respectively. Furthermore, we computed the normalized distance to the cell nuclei centroid. This calculation entailed determining the distance between the query transcript and the nuclei centroid, which was then divided by the maximal distance between all transcripts and the nuclei centroid. Finally, we employed kernel density estimation (KDE) plots to artistically depict the smoothed distributions of these distances.

#### Single cell analysis

Single cell analysis was performed using Scanpy^29^. We started by extracting count matrices from the segmented cells, which were then normalized to a total count of 1000 per cell. Cells with insufficient counts (minimum 10 counts per cell) were removed due to low coverage. Then, log transformation and scaling were applied to the normalized counts. Principal Component Analysis (PCA) was subsequently employed to reduce the dimensionality of the data. Using the resulting PCA components, a neighbor graph was constructed by considering the 10 nearest neighbors for each cell. The UMAP algorithm was applied to obtain a reduced-dimensional representation of the data, which facilitated visualization and exploration. To understand cell identities in the results, we used the “ingest” function in Scanpy to map labels from the reference data (annotations in the original paper in this case) to single cell data generated from benchmark methods, including Watershed, Cellpose, ClusterMap and Baysor. Predicted labels from Bering results were used directly for the comparison with other benchmark methods.

#### Alignment of image signals and spot information

DAPI staining images are usually available for image-based spatial technologies. We get the coordinates of transcript spots and project them onto DAPI images based on the closest pixels. The DAPI image intensity of the corresponding pixels were used as the DAPI staining strength of the spots. DAPI image intensity scale from 0 to 255 and 25 is used as the threshold for low signal pixels.

## Data availability

Raw data of all datasets used in the paper can be found in the Supplementary Table 1. Processed data and pre-trained models are deposited in Figshare (https://figshare.com/ndownloader/articles/23605539/versions/2).

## Code availability

The source code and Python package are freely available at https://github.com/jian-shu-lab/Bering. The analysis performed in this paper can be found at https://github.com/jian-shu-lab/Bering_analysis.

## Acknowledgements

This paper is part of the Human Cell Atlas – www.humancellatlas.org/publications/. We thank Jingyi Ren, Jiahao Huang, Hu Zeng and Zefang Tang for the valuable suggestions of STARmap data analysis. We thank Rongxin Fang for sharing thick tissue MERFISH data. Additionally, we thank 10X for discussing Xenium technology details. This work was supported by NIH New Innovator Award (DP2TR004354), NIH Common Fund (UG3CA275687), NICHD (R00HD096049), Massachusetts Life Science Center, Burroughs Wellcome Fund, Additional Ventures, Massachusetts General Hospital, National Institutes of Health LungMap (HL148865), Digestive Health Center (DK078392), National Cooperative Reprogrammed Cell Research Program (MH104172), Center of Excellence in Molecular Hematology (DK126108).

## Author contributions

K.J., Z.Z and J.S. conceived this project. B.A. and J.S. jointly supervised this work. K.J. and Z.Z designed the model, with feedback from J.T.. K.J. conducted benchmark and analysis, with the input from K.Z., F.V. and C.C.. K.J., Z.Z and J.S. wrote the manuscript, with input from all authors.

---

**Figure S1.**
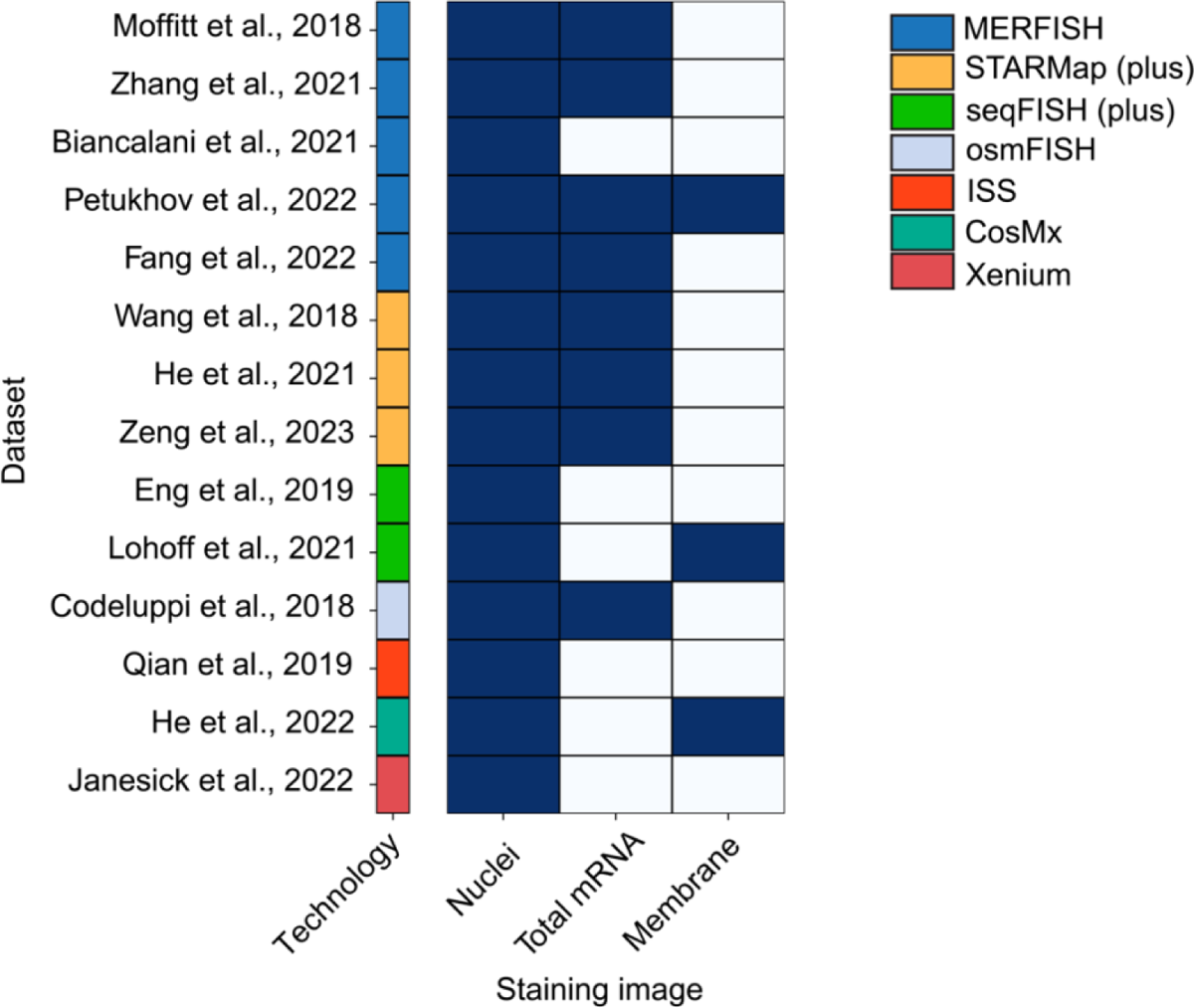
Survey of staining images in various image-based spatial transcriptomics technologies. The table displays the availability of major image channels in different spatial datasets and technologies.

**Figure S2.**
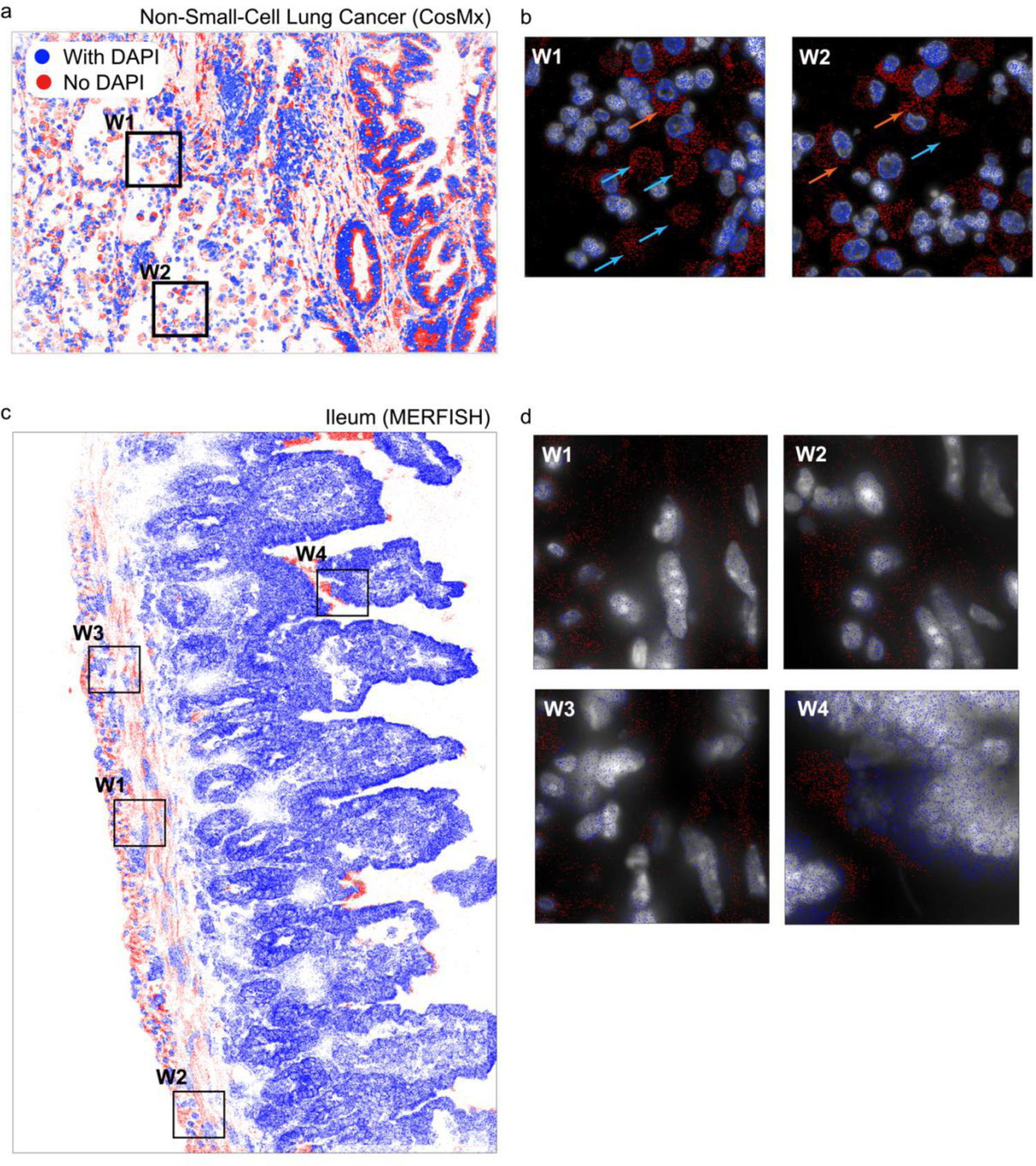
Inadequate image signals in spatial transcriptomics data. (a, c) A CosMx non-small-cell lung cancer slice (a) and a MERFISH mouse ileum slice (c) are presented, where transcripts are visualized as blue and red dots. Blue dots represent transcripts covered by strong DAPI signals, while red dots represent transcripts covered by weak DAPI signals (See Methods). Additionally, selected windows highlight areas with insufficient image staining information. (b, d) Enlarged views of the windows shown in (a) and (c), highlighting DAPI signals in grayscale.

**Figure S3.**
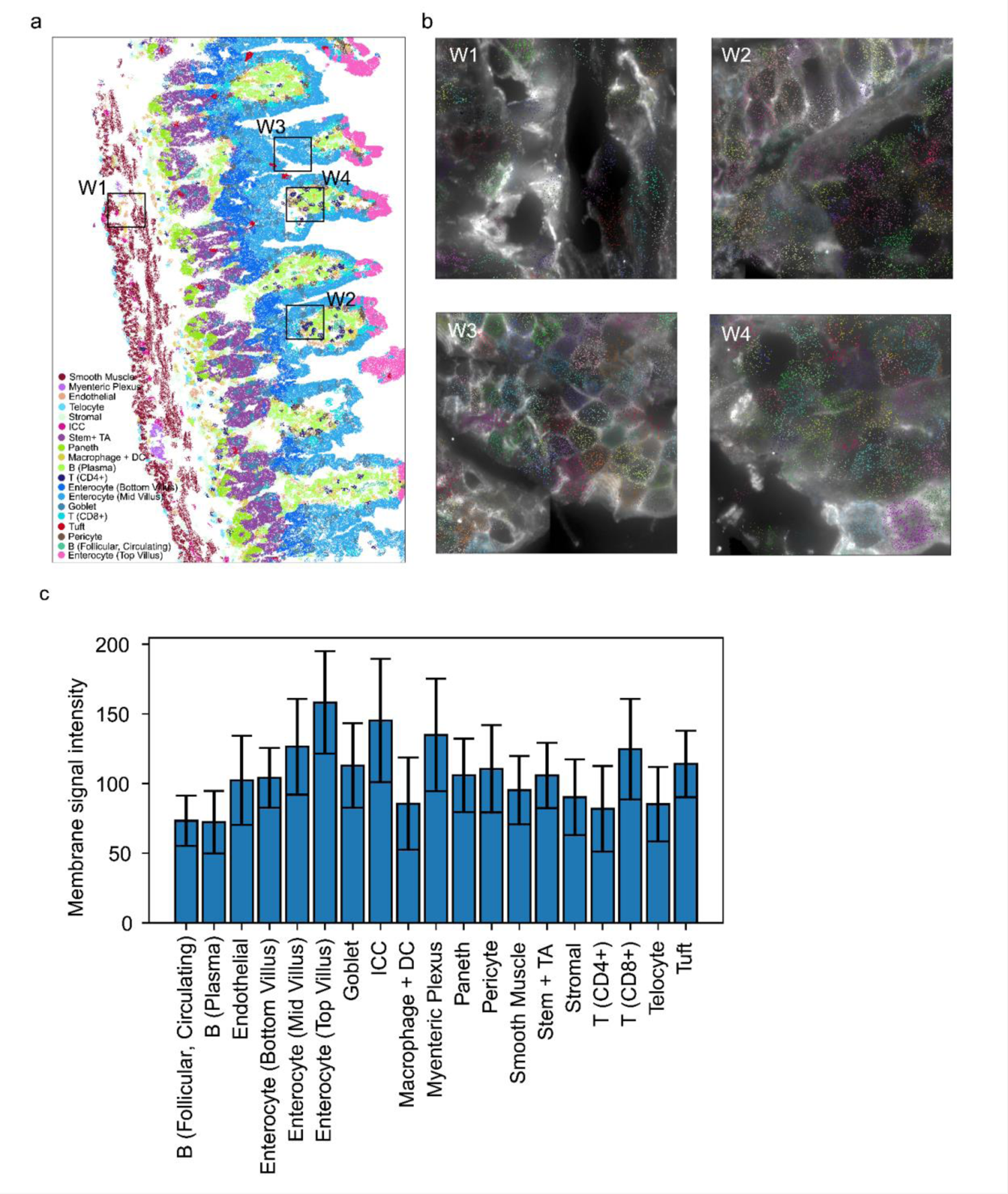
Limitation of cell membrane staining for segmentation. (a) A MERFISH mouse ileum slice is presented, showcasing transcripts from various cell types depicted in distinct colors. Four representative regions are highlighted for further investigation. (b) Enlarged visualization of the four highlighted regions in (a) reveals grayscale membrane imaging, while transcripts in different cells are displayed using unique colors. Inadequate and imbalanced membrane signals are observed across regions, particularly in regions highlighted by W2 and W4. (c) The bar plot illustrates the imbalance in membrane staining signals across different cell types.

**Figure S4.**
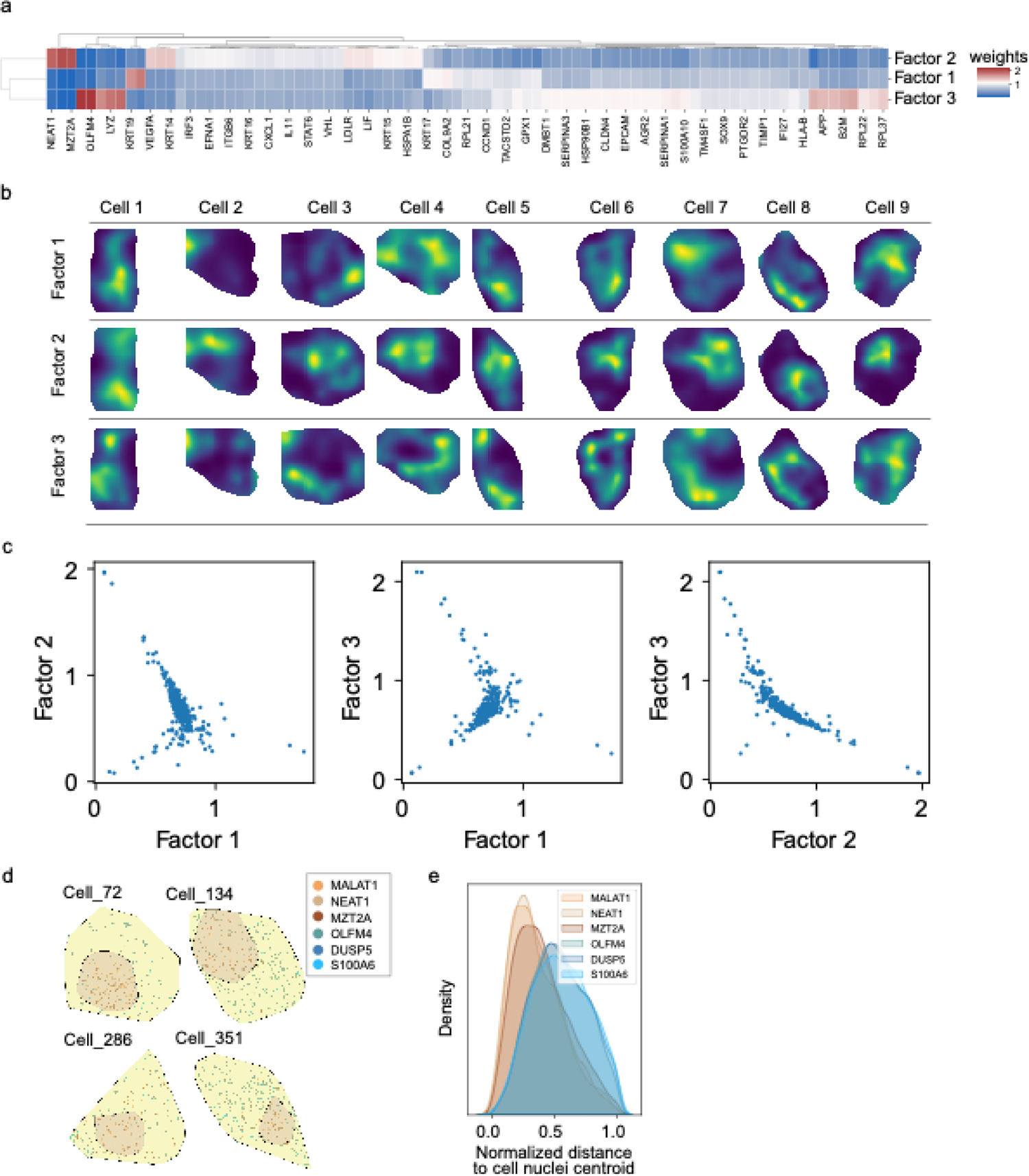
Subcellular patterns of genes in NSCLC tumor cells by FISHFactor. (a) FISHFactor was applied to tumor cells from CosMx NSCLC data, resulting in the identification of three factors representing specific subcellular spatial distributions of genes (see Methods). The weight matrix of genes with high weights across factors is displayed. (b) The visualization of factor scores in nine tumor cells offers valuable insights into the underlying subcellular spatial patterns associated with the factors. Factor 2 reveals high scores primarily in the nuclei region of the cell, suggesting the presence of nuclear genes. Conversely, factor 1 demonstrates high scores at the cell periphery, indicating the potential distribution of cytoplasmic or membrane genes. (c) Scatter plots illustrate the associations between factors. Factor 1 and factor 3 display a positive correlation, while factor 2 and factor 3 demonstrate a negative correlation. (d) Six spatially variable genes were derived from factor 2 and 3. Four representative tumor cells are shown, where cytoplasm and nuclei regions were depicted in yellow and orange, respectively. (e) Density curves depict the distribution of transcripts at varying distances from the nuclei centroid within NSCLC tumor cells. These curves indicate relative positioning of transcripts in relation to the nuclei. The estimation of these density curves was performed using Kernel Density Estimation (KDE).

**Figure S5.**
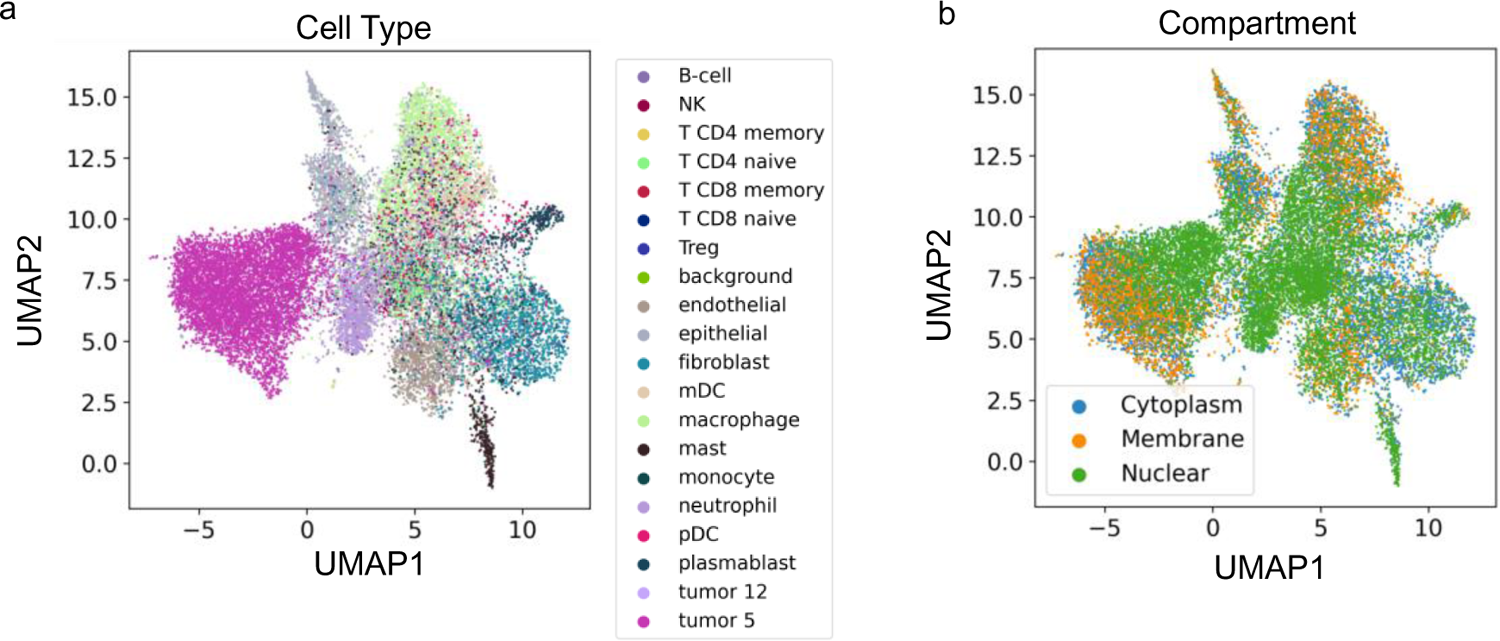
Uniform Manifold Approximation and Projection (UMAPs) depicting NGCs derived from CosMx NSCLC data. Cell type annotations (a) and subcellular compartments (b) for NGCs are displayed in different colors (see Methods).

**Figure S6.**
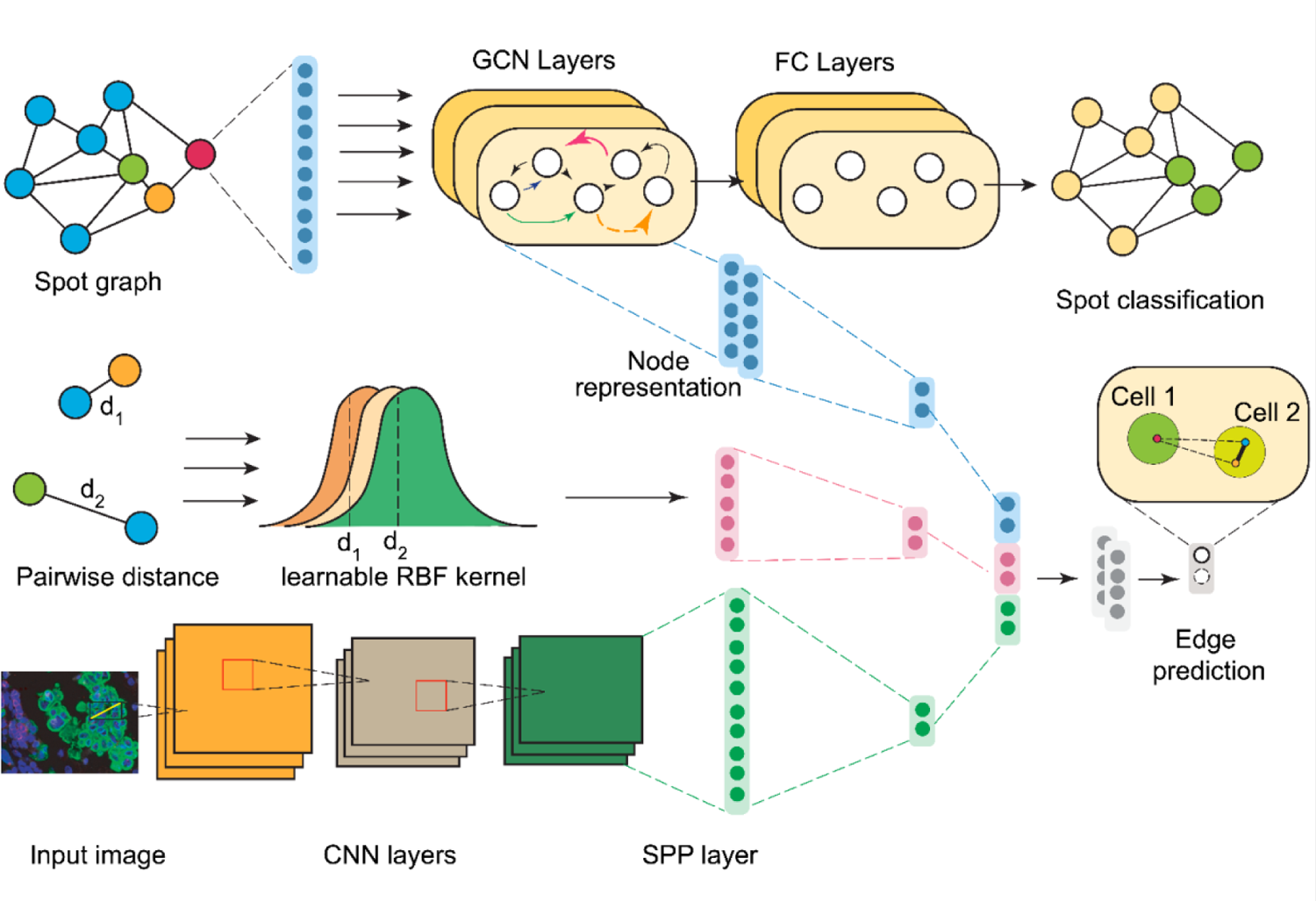
The framework of Bering mode. This framework serves as a supplementary plot to the model framework presented in Fig. 1, providing more detailed information about the neural networks involved. The first part of the Bering model is transcript classification. To accomplish this, a transcript colocalization graph is constructed, followed by the learning of node representations using Graph Convolutional Networks (GCNs). These node representations are utilized for both the transcript classification task and the cell segmentation task, serving as input for edge representation. In addition to node representation, the edge representation incorporates learned distance information and image information. Learnable RBF kernels and Convolutional Neural Networks (CNNs) are employed to capture distance and image features, respectively. The concatenated edge representation is then utilized as input for the edge prediction task, aiming to predict whether two nodes connected by an edge originate from the same cell.

**Figure S7.**
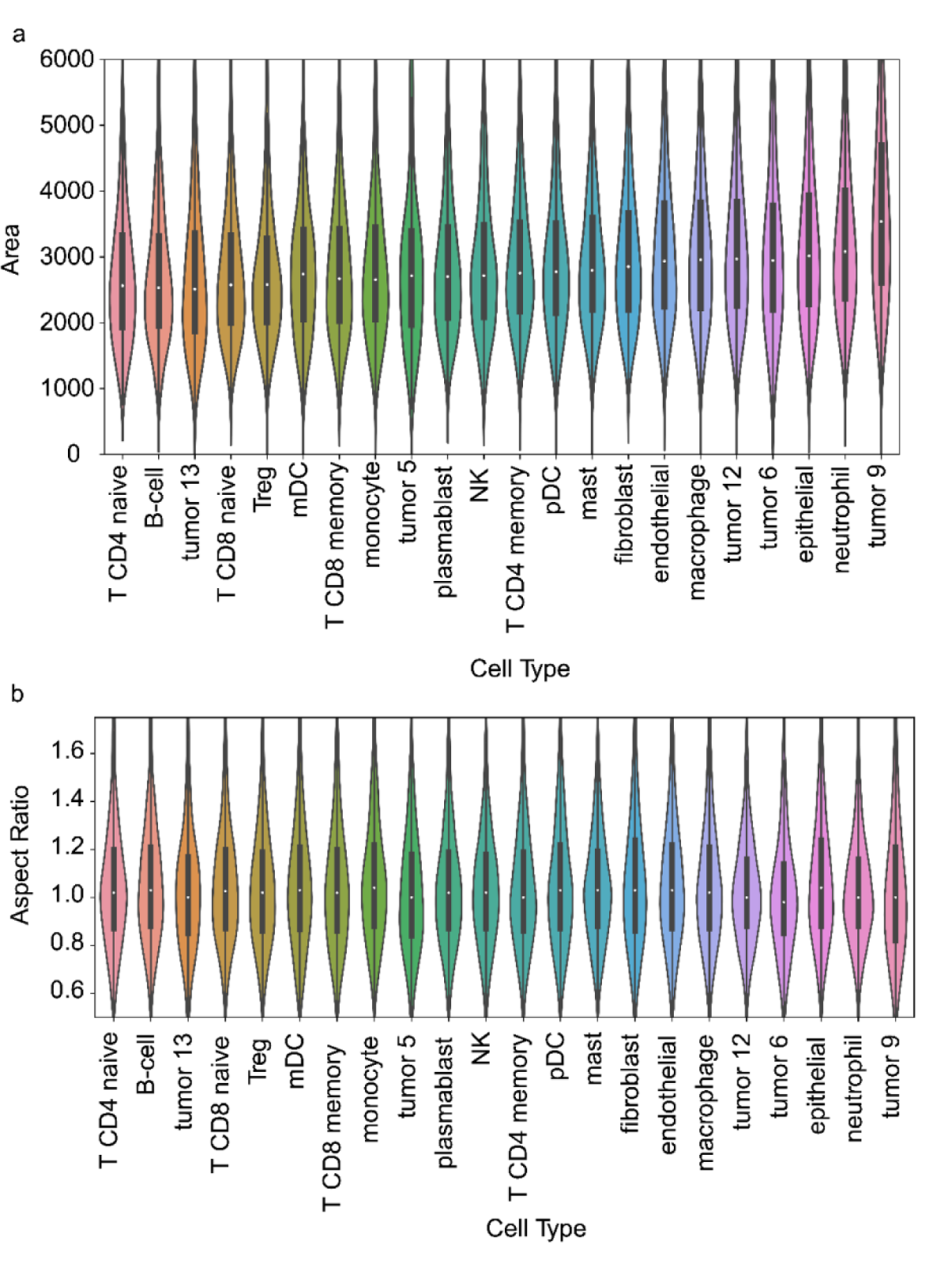
Cell size and shape heterogeneity. (a-b) Violin plots displaying the wide dispersion of cell areas (a) and aspect ratios (b) within specific cell types or across different cell types, using CosMx NSCLC data as an example.

**Figure S8.**
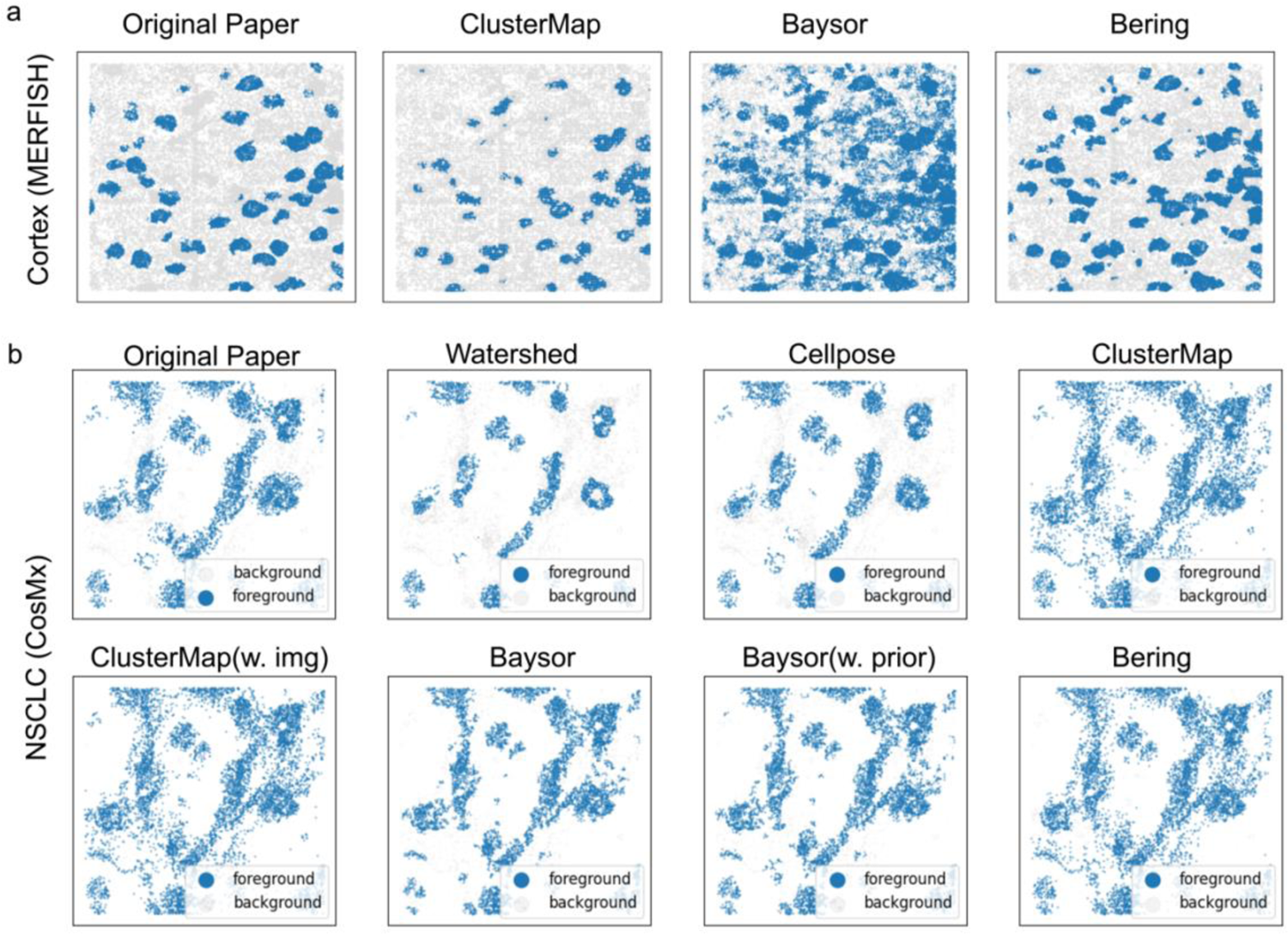
Prediction of real signals and noises in real cases. (a, b) Two windows displaying foreground and background predictions using different segmentation algorithms in the mouse cortex MERFISH data (a) and the non-small cell lung cancer CosMx data (b). Real signals are denoted by blue dots, while background noises are represented by gray dots.

**Figure S9.**
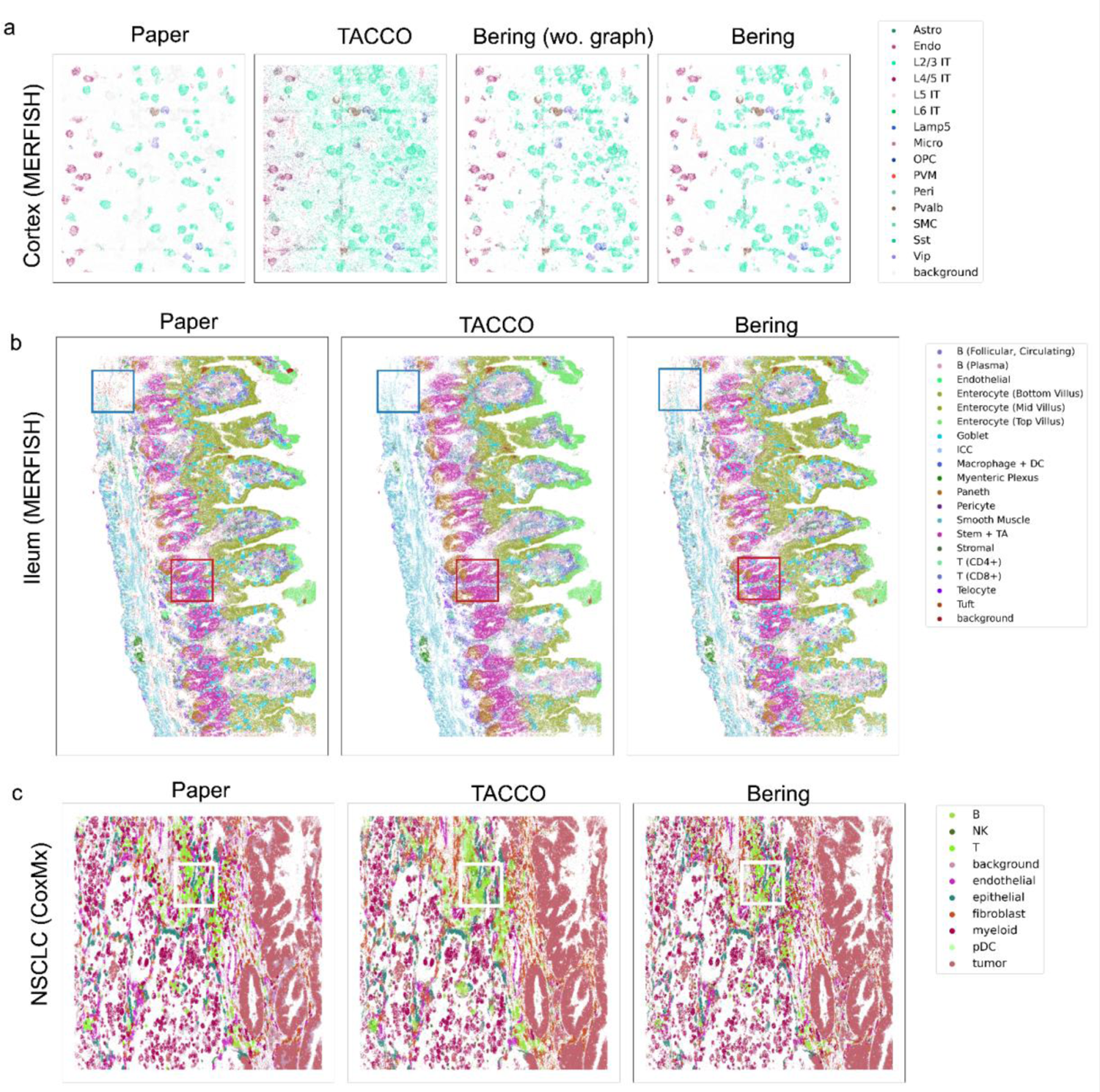
Node classification performance of TACCO and Bering. (a) Two enlarged views of mouse cortex MERIFSH slice showing original labels, TACCO predicted labels, and Bering labels (with and without graph models) from left to right. Different cell types are color-coded, while background noises are depicted in gray. (b-c) Original labels and predicted labels from TACCO and Bering (with graph models) in mouse ileum (b) and NSCLC (c). Highlighted boxes depict microenvironment regions where Bering predicts more details compared with TACCO.

**Figure S10.**
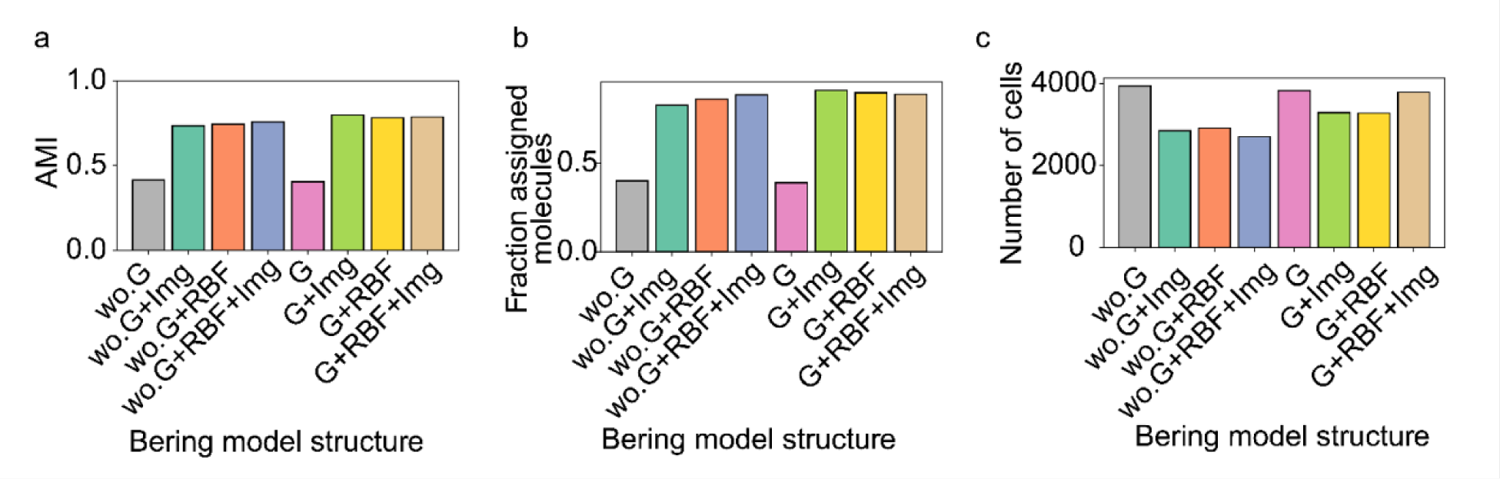
Ablation studies of the Bering model. Ablation studies were conducted to assess the impact of various model components, including graph models, RBF distance kernels, and image embeddings. The segmentation performance was assessed by evaluating different combinations of these model components using quantitative metrics such as Adjusted Mutual Information (AMI), fraction of assigned molecules, and number of detected cells. The analysis was performed on NSCLC CosMx data. wo.G: without graph models (learn the node representation by fully connected layers); G: with graph models (learn the node representation by GCNs); RBF: RBF distance kernels; Img: image embeddings.

**Figure S11.**
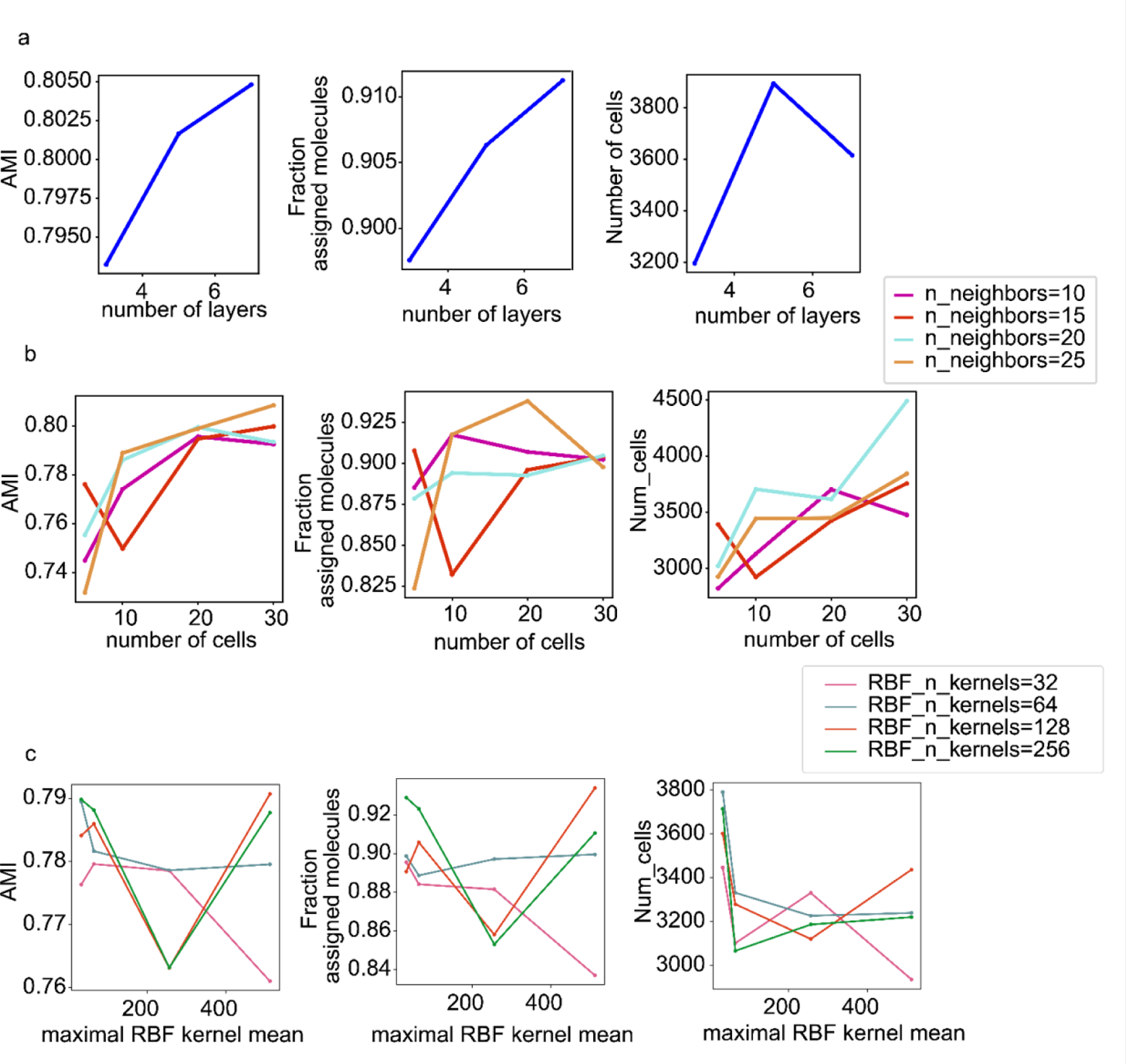
Hyperparameter search of the Bering model. (a-c) The segmentation performance of the Bering model was evaluated in NSCLC data using multiple metrics, including Adjusted Mutual Information (AMI), fraction of assigned molecules, and number of detected cells. Various hyperparameters were examined, including the number of neural network layers in the Bering model (a), the number of neighbors (b), and the number of RBF distance kernels (c).

**Figure S12.**
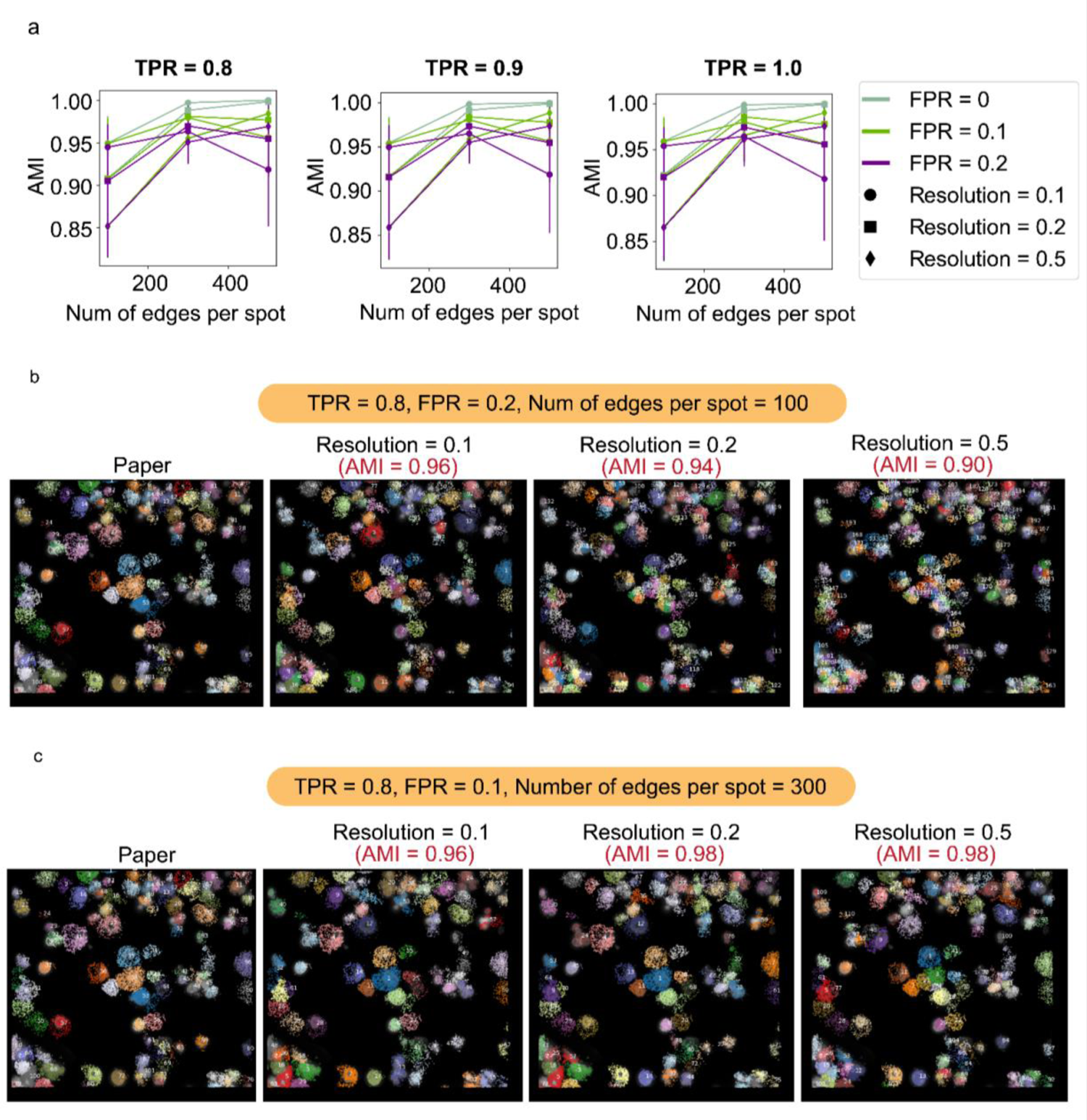
Hyperparameters in the community detection algorithm of cell segmentation. (a) Adjusted Mutual Information (AMI) is utilized to evaluate the segmentation performance across various hyperparameters, including true positive rates (TPR), false positive rate (FPR) in the edge prediction task, and resolutions in Leiden Clustering. Error bars represent standard deviation. (b-c) The community detection performance demonstrates minimal sensitivity to resolution selection when FPR is low in the edge prediction task and there are a higher number of measured edges. Notably, in the edge prediction task with high FPR and low numbers of edges for measurement, the Leiden clustering algorithm shows a reduction in AMI for predicted cells (b). Conversely, it achieves stable and high AMI in the segmentation with low FPR and high numbers of edges for measurement (c).

**Figure S13.**
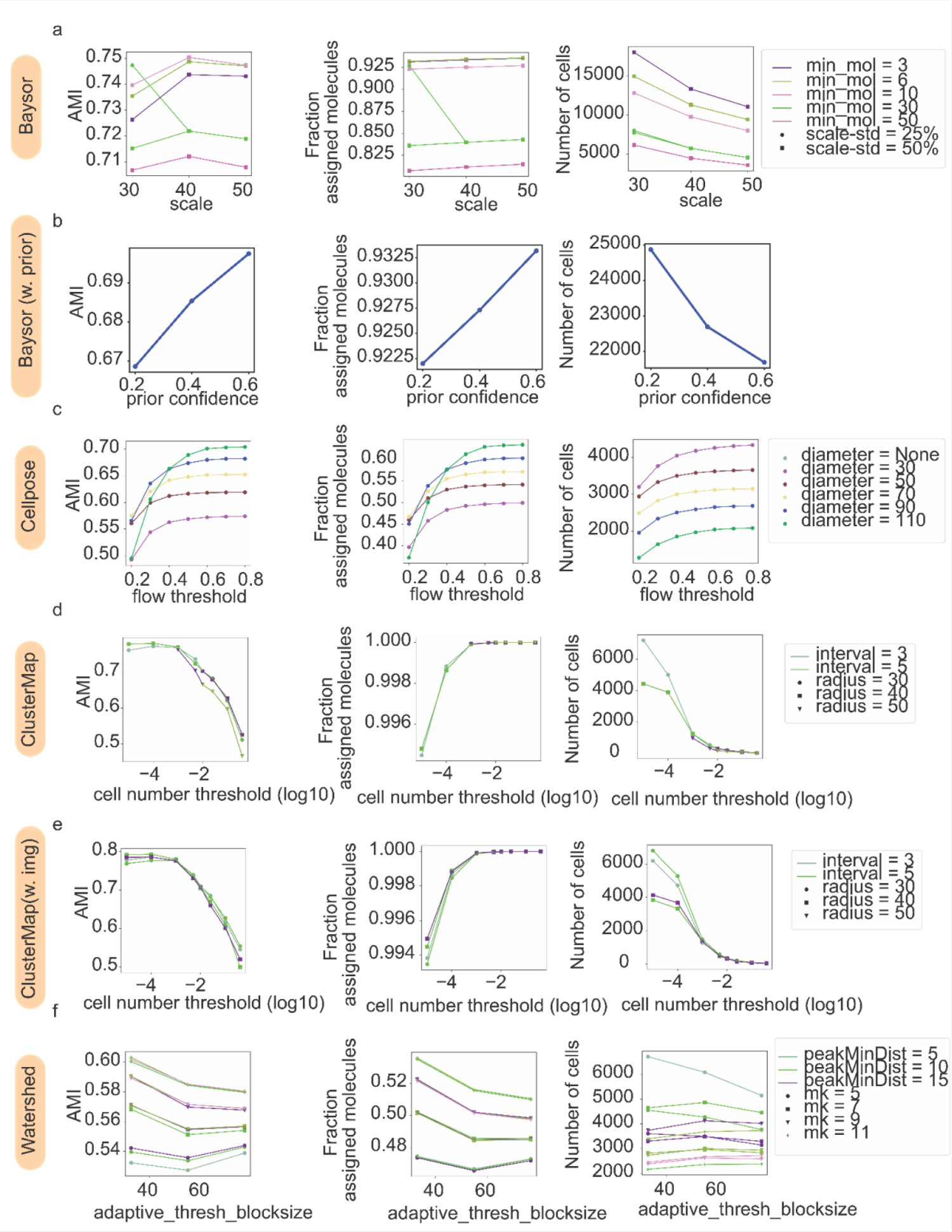
Hyperparameter search of benchmark methods. (a-f) The segmentation performance of the Bering model was evaluated in NSCLC data using multiple metrics, including Adjusted Mutual Information (AMI), fraction of assigned molecules, and number of detected cells. Various hyperparameters were examined for different benchmark approaches, including Baysor (without prior) (a), Baysor (with prior) (b), Cellulose (c), Clustermap (without image) (d), Clustermap (with image) (e), and Watershed (f).

**Figure S14.**
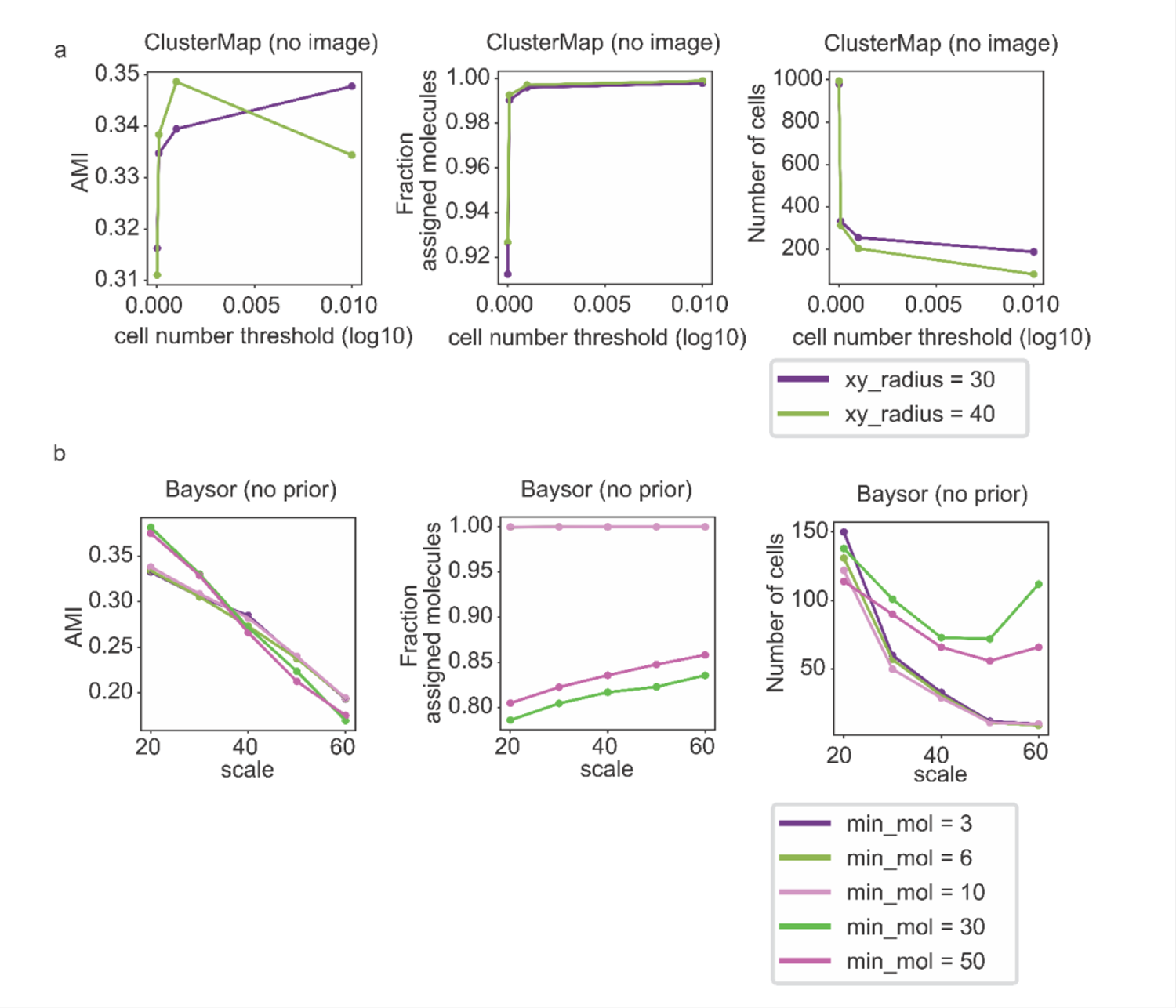
Hyperparameter tuning of benchmark methods for image-free segmentation. (a-b) Image-free segmentation was performed in mouse cortex MERFISH data without the input of images, and various metrics were used to measure the segmentation performance in Clustermap (a) and Baysor (b).

**Figure S15.**
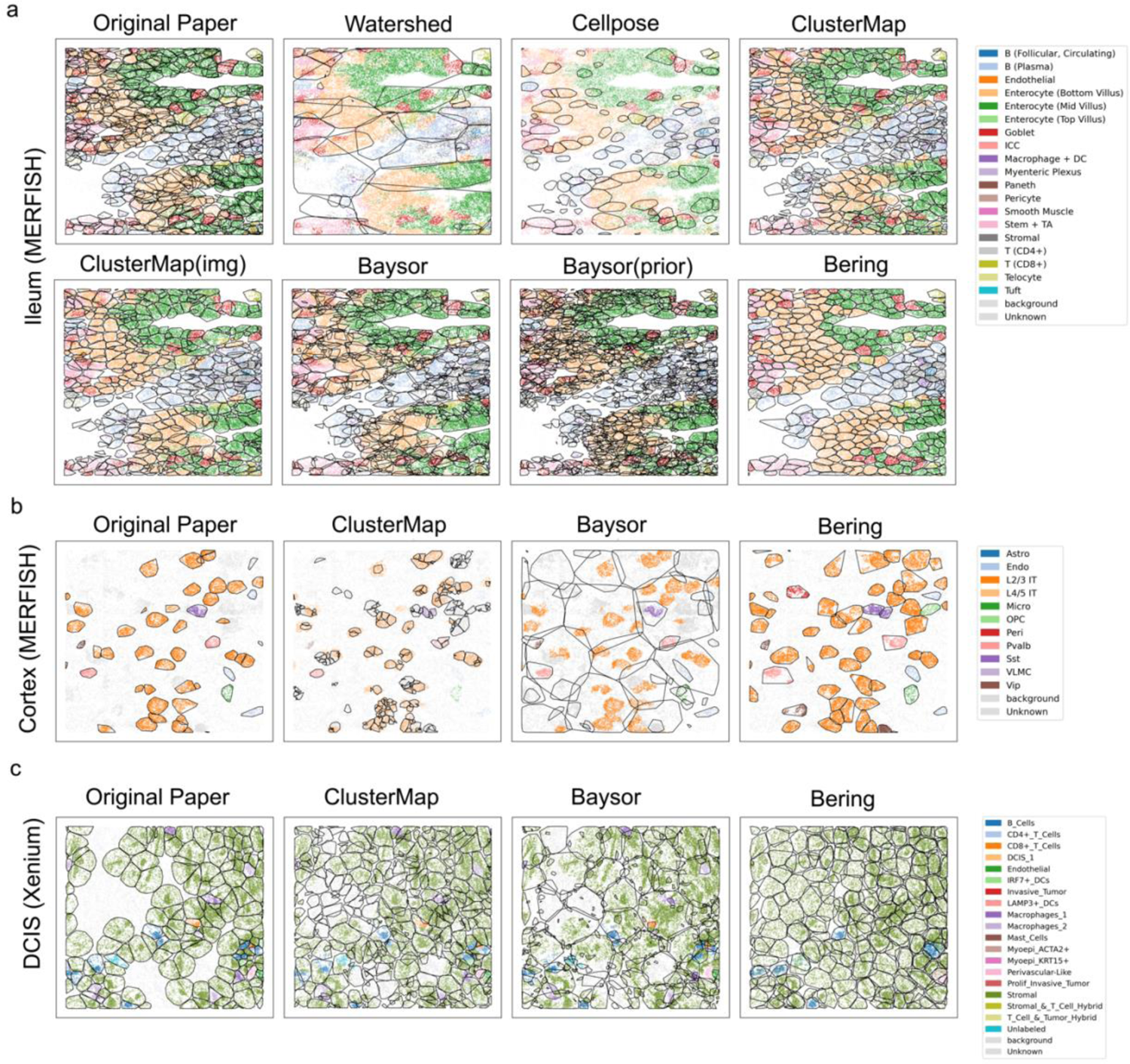
Comparison of cell segmentation methods across datasets is depicted, including Ileum MERFISH data. (a), cortex MERFISH data (b), and DCIS Xenium data (c). In cases where processed nuclei images were unavailable for a dataset, image-dependent segmentation methods were excluded from the comparison.

**Figure S16.**
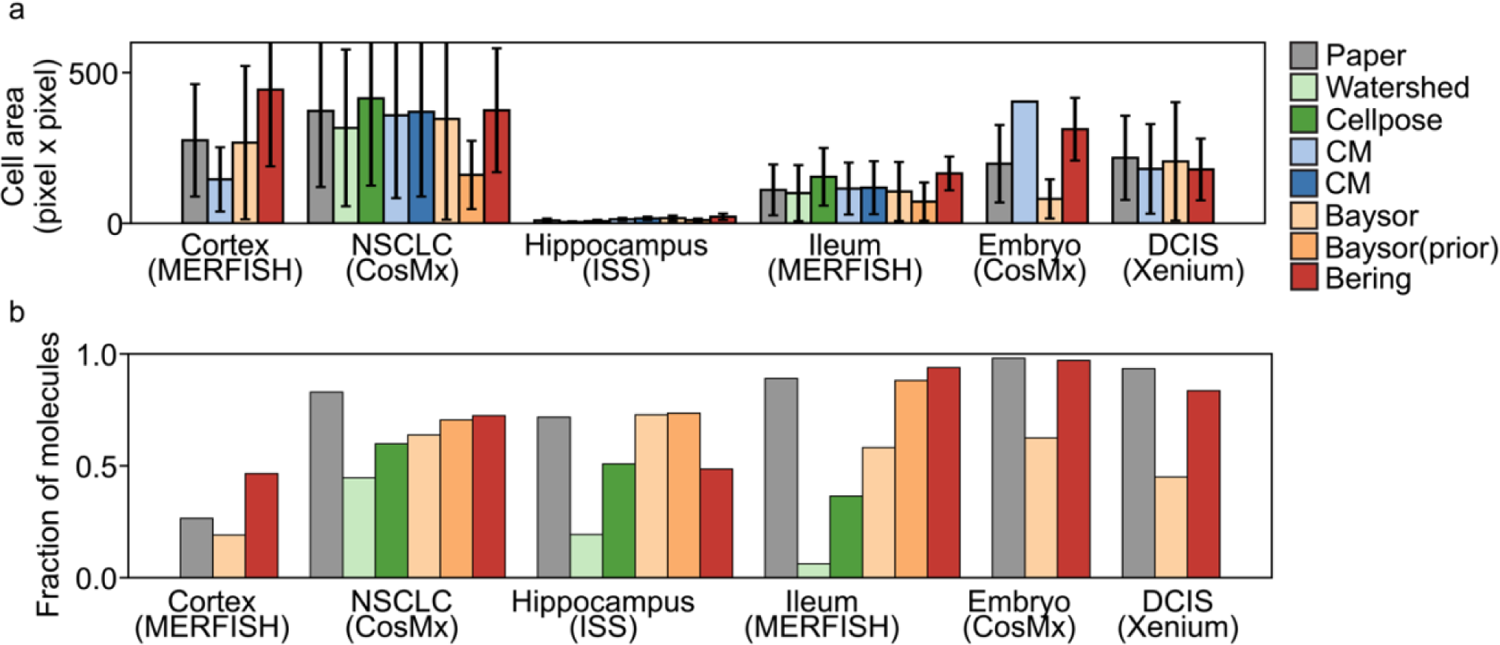
Performance of cell segmentation across datasets. (a-b) Quantitative metrics, such as cell areas (a) and fractions of assigned molecules (b), are employed to benchmark the segmentation results across diverse datasets. The error bars represent the standard deviations of fractions of assigned molecules. Image-dependent methods were excluded from the benchmark if processed nuclei staining images were unavailable.

**Figure S17.**
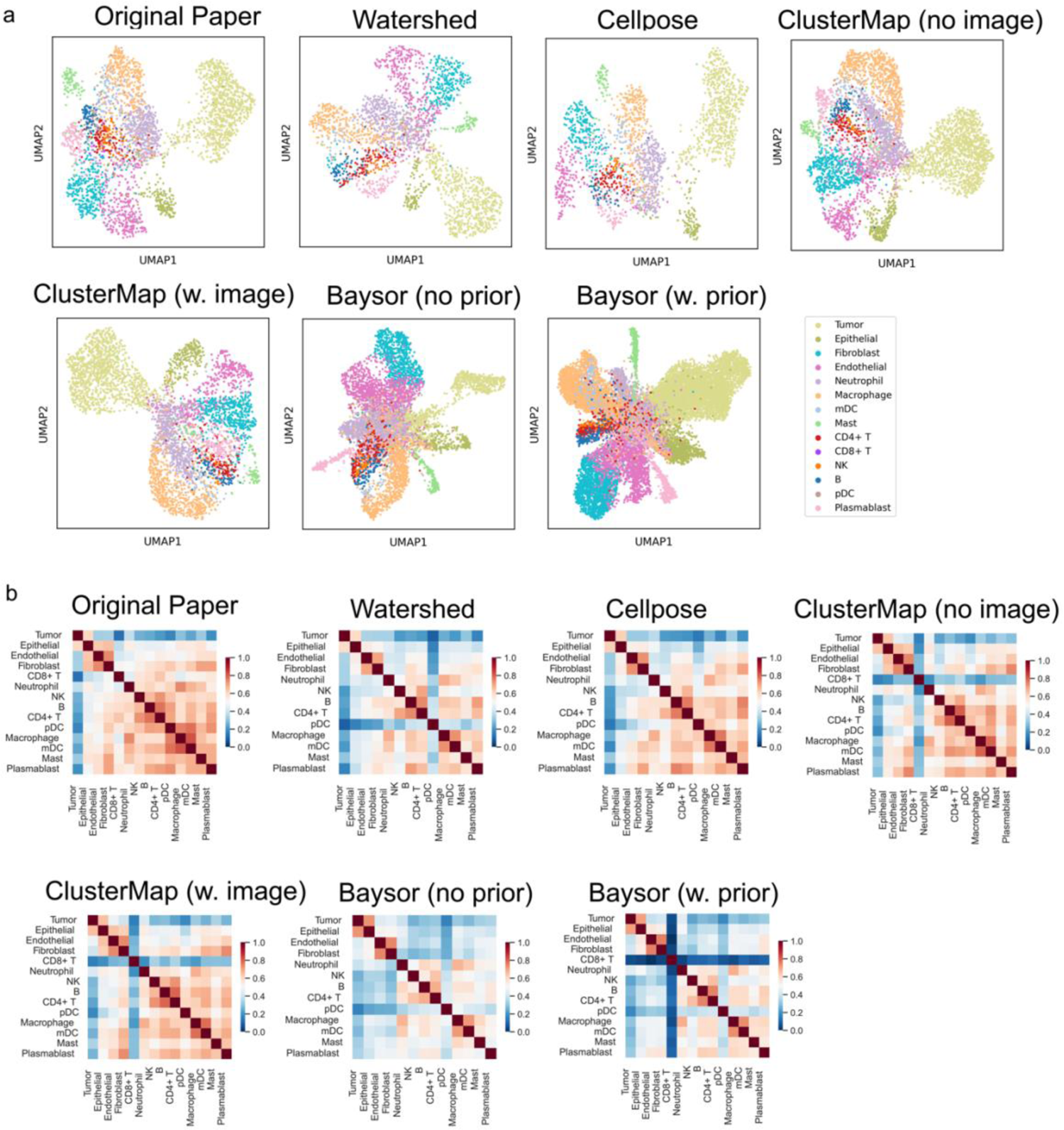
Comparison of single-cell analysis in different benchmark methods. (a) UMAP plots illustrating single-cell clustering results using segmented cells obtained from the original papers (top left) and other benchmark methods (See methods). (b) Spearman correlation computed for gene expression levels in the clusters depicted in (a). Representative cell markers, identified through differential expression analysis from the original single-cell data in the paper, were used for correlation measurement. For Bering results, please refer to Figure 3.

**Figure S18.**
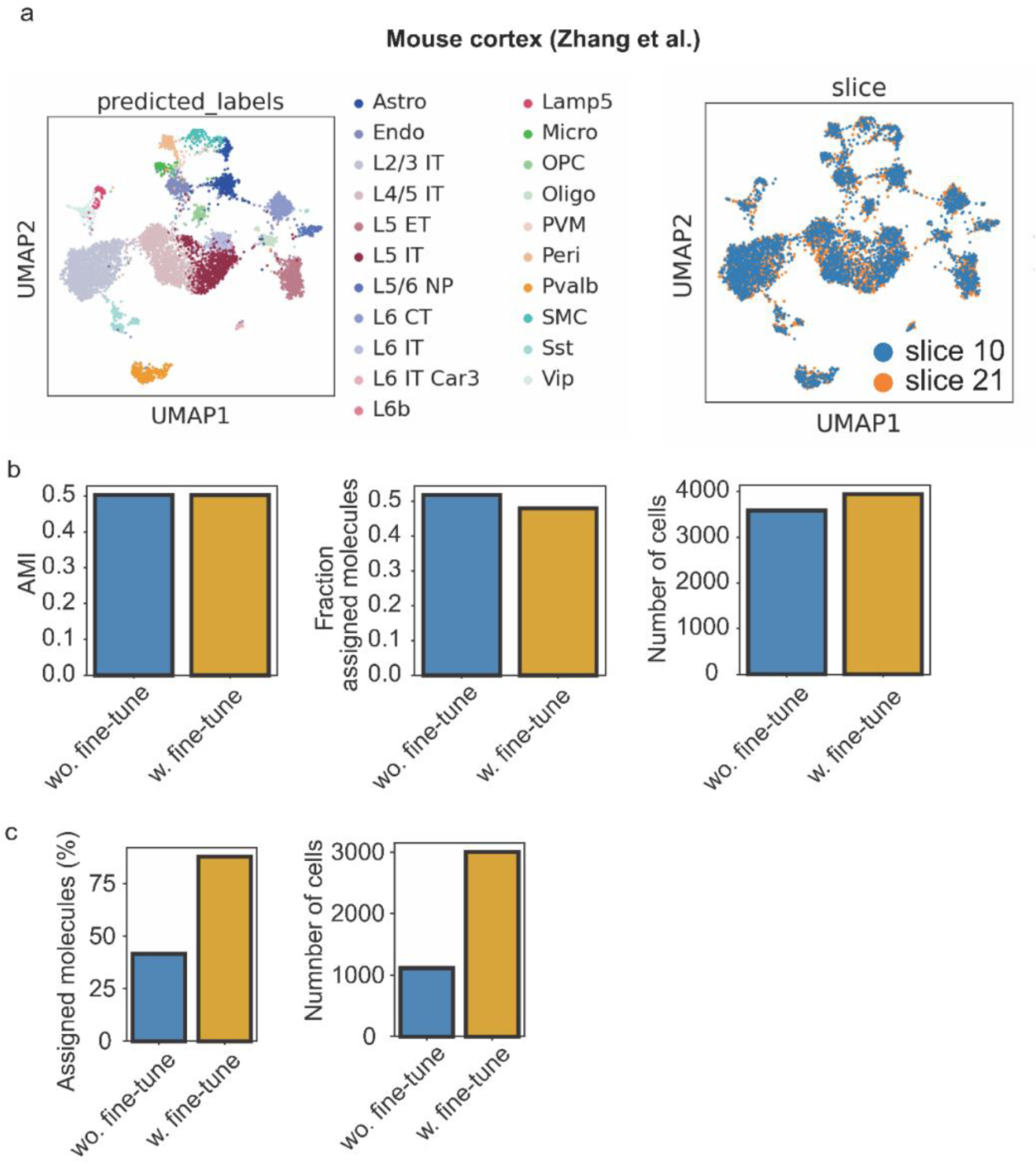
Generalizability of the Bering model within and across datasets. (a) UMAP plots displaying the labels (left) and sources (right) of integrated single-cell data from slice 21 and slice 10 of the MERFISH mouse cortex dataset. The pre-trained model was obtained from slice 21 and applied to slice 10. The cells and labels in slice 10 were derived from the segmentation prediction results. (b) Performance comparison between model application with and without fine-tuning of the pre-trained Bering model for the task described in (a). Quantitative metrics, including adjusted mutual information (AMI), fraction of assigned molecules, and number of segmented cells, were measured for both strategies in slice 10. (c) The segmentation performance of the pre-trained tumor model with and without fine-tuning. The pre-trained model was obtained from NSCLC CosMx data and applied in DCIS Xenium data, as illustrated in Fig. 5e-g.

